# Synthesis and pharmacological characterization of UVI3502, a novel cannabinoid receptor 1 (CB_1_) antagonist/inverse agonist

**DOI:** 10.1101/2024.06.14.598980

**Authors:** Iker Bengoetxea de Tena, Gorka Pereira-Castelo, Jonatan Martínez-Gardeazabal, Marta Moreno-Rodríguez, Iván Manuel, Claudio Martínez, Belén Vaz, Javier González-Ricarte, Rosana Álvarez, Angel Torres-Mozas, Francesca Peccati, Gonzalo Jiménez-Osés, Ángel Rodríguez De Lera, Rafael Rodríguez-Puertas

**Affiliations:** Department of Pharmacology, Faculty of Medicine and Nursing, University of the Basque Country (UPV/EHU), Leioa, Spain; Neurodegenerative Diseases, BioBizkaia Health Research Institute, Barakaldo, Spain; Department of Organic Chemistry, Faculty of Chemistry and CINBIO, University of Vigo, Vigo, Spain; Center for Cooperative Research in Biosciences (CIC BioGUNE), Basque Research and Technology Alliance (BRTA) Bizkaia Technology Park, Derio, Spain; Ikerbasque, Basque Foundation for Science, Bilbao, Spain

## Abstract

The endocannabinoid (eCB) system regulates several brain functions and is implicated in neurological disorders. The pharmacological blockade of cannabinoid receptors has a therapeutic potential for various cognitive deficits, but also produces severe psychiatric side effects. Hence, new cannabinoid compounds that potentiate therapeutic effects, while minimizing toxicity, are required. In this study, we synthesized and characterized a novel antagonist/inverse agonist of CB_1_ receptors. UVI3502 showed affinity for two [^3^H]CP55,940 binding sites (IC_50Hi_ 0.47 ± 1.94 nM and IC_50Lo_ 1470 ± 1.80 nM). Subsequent binding assays performed in CB_1_ and CB_2_ overexpressing membranes determined that the low affinity binding site corresponded to CB_1_, but the high-affinity binding site of UVI3502 did not correspond to CB_2_ and the possibility of it corresponding to GPR55 was analyzed. The affinity of UVI3502 for CB_1_ receptors was further confirmed with neuroanatomical specificity by autoradiography in key brain areas, in which functional [^35^S]GTPɣS assays demonstrated that UVI3502 behaved as an antagonist/inverse agonist of CB_1_ receptors, blocking the stimulation evoked by potent cannabinoid receptor agonist CP55,940 and decreasing basal [^35^S]GTPɣS binding. The *in silico* characterization of the binding to CB_1_ receptor through molecular docking and molecular dynamics suggests that this activity is explained by the planar and rigid structure of UVI3502, which is optimal for interactions with the inactive state of the receptor. These results indicate that UVI3502 is a novel antagonist/inverse agonist of CB_1_ receptors, making it a compelling candidate for pharmacologically blocking cannabinoid receptors in the central nervous system.

**Significance Statement:** UVI3502 is a novel antagonist/inverse agonist of CB_1_ receptors, with almost no affinity for CB_2_ receptors and an additional high-affinity binding site for a third, cannabinoid-like receptor, potentially GPR55. In relevant brain areas for learning and memory processes with a high expression of CB_1_, UVI3502 blocks the stimulation evoked by the cannabinoid receptor agonist CP55,940, rendering it as an interesting compound for the pharmacological blockade of cannabinoid receptors in the central nervous system.

## Introduction

The endocannabinoid (eCB) system is essential for preserving energy balance, metabolism, and energy expenditure (Cristino et al., 2014), but is also implicated in modulating cognitive functions, including learning and memory (Lu & Mackie, 2021). This system involves two identified G protein-coupled receptors (GPCRs): the type-1 and -2 cannabinoid receptors, CB_1_ and CB_2_. Their expression differs significantly, with CB_1_ being widely expressed in the central nervous system (CNS), including in brain areas associated with the psychoactive effects of Δ9-tetrahidrocannabinol (Δ9-THC); the basal ganglia, the cerebellum, portions of the hippocampus and the cortical regions (Mackie, 2008). In contrast, CB_2_ receptors are primarily expressed in the immune system and the spleen (Berdyshev, 2000; Schatz et al., 1997). Besides CB_1_ and CB_2_, additional GPCRs showing little amino acid sequence homology have also been described as “cannabinoid-like” receptors, due to their affinity for endocannabinoids and/or synthetic cannabinergic drugs. Among them, GPR55 receptors are the most notable example and are of particular interest due to the relatively high levels of expression in the brain, particularly in the striatum (Sawzdargo et al., 1999).

As one of the most relevant neuromodulatory networks of the CNS, the eCB system regulates important physiological processes, such as neurodevelopment, synaptic plasticity and adaptive responses (Cristino et al., 2020; Lu & Mackie, 2021), all of which affect cognition. Hence, in spite of the well-known deleterious effects of cannabinoids on memory (Broyd et al., 2016), there is a growing interest in developing new cannabinoid compounds for the treatment of neurological and neurodegenerative diseases (Arachchige, 2023; Voicu et al., 2023). A prime example of this are Alzheimer’s disease (AD) and related dementias, given the very limited efficacy of the currently approved drugs for these disorders (Gharat et al., 2024). In this context, various components of the eCB system undergo alterations in *postmortem* human samples from AD patients (Manuel et al., 2014; Moreno-Rodriguez et al., 2024; Westlake et al., 1994). Importantly, preclinical evidence using animal models of AD also suggest that pharmacological or genetic manipulations of the eCB system affect cognition and can influence the histopathological and biochemical markers associated with this disease (Aso et al., 2012; Llorente-Ovejero et al., 2018). In addition to AD, preclinical studies also show the potential of cannabinoid treatments in animal models of other neurodegenerative disorders frequently linked with dementia, such as Parkinson’s disease (PD) and Huntington’s disease (HD) (Mounsey et al., 2015; Sagredo et al., 2009). Besides neurodegeneration, in a mouse model of Williams-Beuren syndrome, a neurodevelopmental disorder characterized by social and cognitive abnormalities, an indirect and subchronic stimulation of the eCB system restored these deficits (Navarro-Romero et al., 2022).

While most of these treatments involve the activation of the eCB system, either through direct action on cannabinoid receptors or through the modulation of the synthesis and degradation enzymes of endocannabinoids, the pharmacological blockade of cannabinoid receptors also exerts positive effects depending on the context. For instance, in two mouse models of Down’s syndrome (DS), treatments with the gold-standard antagonist/inverse agonist of CB_1_ receptors, SR141716A or rimonabant (Rinaldi-Carmona et al., 1995), restored key cognitive phenotypes affected by the pathology (Navarro-Romero et al., 2019). Similarly, in another neurodevelopmental condition, fragile X syndrome, CB_1_ blockade normalized cognitive impairment and altered spine morphology, whereas a pharmacological blockade of CB_2_ only normalized anxiolytic-like behavior (Busquets-Garcia et al., 2013; Gomis-González et al., 2016). In a very different context, mice treated with the well-known muscarinic antagonist scopolamine, known to produce transient cholinergic hypofunction and cognitive deficits, co-treatment with MK-7128, a CB_1_ receptor inverse agonist, improved performance in different behavioral tasks and this was achieved at moderate levels of CB_1_ occupancy in the brain (Dillon et al., 2011).

With this in mind, the pharmacological inhibition of cannabinoid receptors, particularly CB_1_, offers the potential to ameliorate a wide arrange of cognitive deficits arising from various causes. However, the most used compound in these studies, rimonabant, is a very high affinity inverse agonist which produced severe psychiatric side effects, including anxious and depressive disorders, when it was administered to patients suffering from obesity (Moreira & Crippa, 2009; Pi-Sunyer et al., 2006). Thus, there is a need to develop new compounds with a similar pharmacological profile that avoid such adverse effects.

We have therefore synthesized and characterized, both *in vitro* and *in silico*, a novel and subtype specific antagonist/inverse agonist of CB_1_ receptors, UVI3502, which blocks the stimulation evoked by potent cannabinoid agonist CP55,940 in some of the most relevant brain areas for learning and memory processes.

## Materials and Methods

### Reagents, drugs and chemicals

All the compounds necessary for the different procedures were of the highest commercially available quality for the purpose of our studies. For the synthesis of the novel compounds, chemical reagents of the highest purity available were purchased from Sigma-Aldrich and used as received, except when indicated.

[^3^H]CP55,940 (149 Ci/mmol) and [^35^S]GTPɣS (1250 Ci/mmol) were acquired from Revvity (Waltham, MA, USA). The [^3^H]-microscales and [^14^C]-microscales used as standards in the autoradiographic experiments were purchased from ARC (American Radiolabeled Chemicals, Saint Louis, MO, USA). The β-radiation sensitive films, Kodak Biomax MR, bovine serum albumin (BSA), DL-dithiothreitol (DTT), guanosine 5’-diphosphate (GDP), guanosine 5’-O-3-thiotriphosphate (GTPɣS), ketamine and xylazine were all acquired from Sigma-Aldrich (St Louis, MO, USA).

(-)-cis-3-[2-Hydroxy-4-(1,1-dimethylheptyl)phenyl]-trans-4-(3-hydroxypropyl) cyclohexanol (CP55,940), 5-(4-Chlorophenyl)-1-(2,4-dichlorophenyl)-4-methyl-N-1-piperidinyl-1H-pyrazole-3-carboxamide hydrochloride (SR141716A) and (11R)-2-Methyl-11-[(morpholin-4-yl)methyl]-3-(naphthalene-1-carbonyl)-9-oxa-1-azatricyclo[6.3.1.04,12]dodeca-2,4(12),5,7-tetraene (WIN55,212-2) were acquired from Tocris (Bristol, UK). [(1S,2S,5S)-2-[2,6-Dimethoxy-4-(2-methyloctan-2-yl)phenyl]-7,7-dimethyl-4-bicyclo[3.1.1]hept-3-enyl]methanol (HU308) and [(2R)-2-hydroxy-3-[hydroxy-[(2R,3R,5S,6R)-2,3,4,5,6-pentahydroxycyclohexyl]-oxyphosphoryl]oxypropyl] hexadecanoate (lysophosphatidylinositol, LPI) were acquired from Merck (Darmstadt, Germany).

### Chemical synthesis of the novel compounds

We have previously described the palladium-catalyzed heterocyclization/oxidative Heck coupling cascade as synthetic approach to fused heterocycles, including 3-alkenyl-substituted benzofurans, indoles, 1*H*-isochromen-1-imines, tetrahydrodibenzofurans and tetrahydrobenzo[*c*]chromen-6-imines (Álvarez et al., 2010; Álvarez et al., 2012; Martínez et al., 2009; Martínez et al., 2012; Vaz et al., 2021) and extended the procedure to the synthesis of analogues with 7,12-dihydroindolo[3,2-d]benzazepine-6(5*H*)-one skeleton. The synthetic protocol allowed the regioselective construction of the core indole and benzazepinone heterocycles of polycyclic compounds, also known as alkylidenepaullones, which were further characterized as activators of the epigenetic enzymes NAD^+^-dependent class of histone deacetylases (sirtuins, Sirt1) in biochemical assays (Denis et al., 2015). The complete information regarding the synthesis of each of the novel compounds is described in detail in the Supplemental Data.

### Animals, tissues and cells

Every effort was made to minimize animal suffering and to use the minimum number of animals possible throughout the whole study. All procedures using all animal species were performed in accordance with the Guide for the Care and Use of Laboratory Animals as adopted and promulgated by the U.S. National Institutes of Health, with the European animal research laws (Directive 2010/63/EU) and the Spanish National protocols, and were approved by the Local Ethical Committee for Animal Research of the University of the Basque Country (CEEA M20-2018-52 and 54).

### Sprague-Dawley rats

Male Sprague-Dawley rats weighing 200-300 g were housed in groups of 3-4 per cage at a temperature of 22°C and in a humidity-controlled (65%) room with a 12:12 hours light/dark cycle, with access to food and water *ad libitum*. Spleens and brains from Sprague-Dawley (n = 5) rats were used to prepare membrane homogenates for radioligand affinity assays. Brains from Sprague-Dawley rats (n = 5) were also used for autoradiographic studies.

### Swiss mice

Male Swiss mice weighing 20-30 g were housed in a single cage at a temperature of 22°C and in a humidity-controlled (65%) room with a 12:12 hours light/dark cycle, with access to food and water *ad libitum*. Brains from control Swiss mice (n = 5) were used for autoradiographic studies.

### Membrane homogenates from CHO cells overexpressing CB_1_ and CB_2_ receptors

Membrane homogenates from CHO cells overexpressing CB_1_ and CB_2_ receptors, as well as matched wild-type cells, were used to test newly synthesized compounds and were acquired from Sigma-Aldrich (St Louis, MO, USA).

### Preparation of membrane homogenates

Sprague-Dawley rats (n = 5) were anesthetized and sacrificed by decapitation. Spleens and brains were then quickly removed by dissection at 4°C and, in the case of brain tissue, the cortex was dissected for the preparation of the membrane homogenates. For this procedure, spleen and cortex samples were homogenized using a Teflon-glass grinder (15 up-and-down strokes at 800 rpm) in 30 volumes of homogenization buffer (1 mM EGTA, 3 mM MgCl_2_, 50 mM Tris-HCl; pH 7.4) supplemented with 0.25 mM sucrose, at 4°C. The obtained homogenates were centrifuged for 5 min at 1,500rpm. Pellets were removed and supernatants were centrifuged again for 15 min at 14,000 rpm. For washing, the obtained pellets were resuspended in a buffer, centrifuged again and the supernatant was removed. The resulting aliquots were stored at -80°C until use.

### Radioligand binding assays

#### [^3^H]CP55,940 binding assays

To screen the affinity of the newly synthesized compounds for cannabinoid receptors, these were used in concentrations ranging from 10^–12^ M to 10^–4^ M and incubated with a protein concentration of 0.1 mg/mL of rat cortex homogenates for 2 h at 37 °C with agitation in the presence of 0.5 nM of [^3^H]CP55,940. Nonspecific binding was defined as the binding of [^3^H]CP55,940 in the presence of 10^–4^ M of SR141716A. After incubation, the reaction was stopped by adding an ice-cold wash buffer (Tris-HCl 50 mM and 0.5% of BSA, pH 7.4). Then, the membranes were retained by vacuum filtration to a Whatman GF/C glass microfiber filter (Sigma-Aldrich, St. Louis, MO, USA) and the free radioligand was discarded. Filters with the bound radioligand were transferred to vials containing 5 mL of Ultima Gold cocktail (PerkinElmer, Boston, MA, USA) and measured with a Packard Tri-Carb 2200CA liquid scintillation counter (PerkinElmer, Boston, MA, USA).

After that, the compound with the best affinity was tested in cell membrane homogenates overexpressing CB_1_ and CB_2_ receptors. A concentration of 0.02 mg/mL of commercial WT (as control), CB_1_ and CB_2_ overexpressing CHO cells were used, and the same protocol described above was followed. Rat spleen homogenates were also used at a concentration of 0.1 mg/mL as a further characterization of binding to CB_2_ receptor, given the high expression of this receptor in this tissue and the practical lack of CB_1_ in it (Pertwee, 1997). A binding assay using [^3^H]CP55,940 and LPI was also performed. Nonspecific binding was defined as the binding of [^3^H]CP55,940 in the presence of WIN55,212-2 (CB_1_/CB_2_ agonist), SR141716A (specific CB_1_ antagonist/inverse agonist) or HU308 (specific CB_2_ agonist), depending on the tissue used for each assay.

#### [^3^H]CP55,940 receptor autoradiography

For the performance of cannabinoid receptors autoradiography using [^3^H]CP55,940, fresh frozen sections from brain samples from wild-type Sprague-Dawley rats (n = 5) and Swiss mice (n = 5) were used to test the newly synthesized compound which had shown the best affinity for cannabinoid receptors.

All brain sections were air dried for 30 min and then immersed in Coplin jars for preincubation in a buffer containing 50 mM Tris-HCl and 1% of BSA (pH 7.4) for 30 min, at room temperature, to remove endogenous ligands. Two tissue slices were later incubated in the presence of the [^3^H]CP55,940 radioligand (3 nM) for 2 h at 37°C and, in two consecutive slices, the incubation was performed also in the presence of the target compound (10 µM) and in the presence of the known CB_1_ antagonist/inverse agonist SR141716A (10 μM). Following incubation, tissue slices were washed with an ice-cold preincubation buffer, dipped in distilled water and dried overnight. To generate autoradiograms, dry sections were exposed to β-radiation-sensitive films in hermetically closed cassettes for 21 days at 4°C. For the calibration of the optical densities to fmol/mg tissue equivalent, [^3^H]-microscales were exposed to the films. Calibrated films were scanned and quantified using Fiji software (Fiji, Bethesda, MA, USA).

#### [^35^S]GTPɣS functional binding assays

The compound with the best affinity was also tested using a functional [^35^S]GTPɣS binding assay to characterize its activity as an agonist or antagonist/inverse agonist. A protein concentration of 0.1 mg/mL of rat cortex homogenates was used for this assay, suspended in a reaction buffer (Tris-HCl 50 mM, EGTA 1mM, MgCl_2_ 3 mM, NaCl 100 mM, 0.5% of BSA; pH 7.4). The target compound was used in concentrations ranging from 10^–11^ M to 10^–4^ M and incubated for 2 h at 37 °C with agitation in the presence of 0.5 nM of [^35^S]GTPɣS and 50 µM of GDP. Basal coupling of [^35^S]GTPɣS to G_i/o_ proteins was determined by incubating the membrane aliquots with the radioligand in the absence of the target compound. Nonspecific binding was defined as the remaining [^35^S]GTPɣS binding in the presence of 10 mM of unlabelled GTPɣS. After the incubation, the same procedure detailed for the [^3^H]CP55,940 binding assay was followed.

#### Functional [^35^S]GTPɣS autoradiography

For the performance of functional autoradiography of cannabinoid receptors, fresh frozen sections from brain samples from wild-type Sprague-Dawley rats (n = 5) and Swiss mice (n = 5) were used to test the newly synthesized compound which had shown the best affinity for cannabinoid receptors.

All brain sections were air dried for 30 min and then immersed in Copling jars for preincubation (4 times, 15 min each time) in an HEPES-based buffer (50 mM HEPES, 100 mM NaCl, 3 mM MgCl_2_, 0.2 mM EGTA, 0.5% BSA; pH 7.4) at 30°C, to remove endogenous ligands. Slices were then incubated for 2 h at 30°C in the same buffer supplemented with 2 mM of GDP, 1 mM of DTT and 0.04 nM of [^35^S]GTPɣS and the target compound (10 µM) alone, as well as the target compound and CP55,940 (10 µM) together. Basal [^35^S]GTPɣS binding was determined in two consecutive slices in the absence of the agonists. Non-specific binding was defined by competition with non-radioactive GTPɣS (10 µM) in another section. Following incubation, slices were washed twice in ice-cold 50 mM HEPES buffer (pH 7.4), dried and exposed for 48 h to β-radiation sensitive film with a set of [^14^C] standards calibrated for [^35^S]. Calibrated films were scanned and quantified using Fiji software (Fiji, Bethesda, MA, USA). The signal obtained for non-specific binding was subtracted previously from both basal and agonist-stimulated binding. Then, the data was expressed as the percentage of stimulation over basal according to the following formula: ([^35^S]GTPɣS agonist-stimulated binding) × 100/([^35^S]GTPɣS basal binding)-100.

### Molecular docking simulations

Molecular docking calculations were performed using GOLD (CCDC Discovery 2020) and the ChemScore fitness function (Jones et al., 1997; Verdonk et al., 2003). The structure of human cannabinoid receptor CB_1_ was taken from PDB 5TGZ (Hua et al., 2016a). The receptor was prepared from docking using UCSF Chimera to add hydrogen atoms (Pettersen et al., 2004). The docking cavity was centered on the α-carbon of Ser263 and allowed to extend in a spherical surrounding region with a 15 Å radius. UVI3502 coordinates were optimized with Gaussian 16 using the ωB97X-D functional (Chai & Head-Gordon, 2008) and 6-31G(d) basis set. The number of genetic algorithm runs was set to 20. Flexible ligand docking was performed allowing ligand torsions around rotatable bonds and keeping receptor coordinates frozen to crystallographic values.

### Molecular dynamics simulations

MD simulations were carried out with Amber 22 using force fields *ff19SB* (receptor) (Tian et al., 2020), GAFF2 (ligand) (He et al., 2020), and OPC (water) (Izadi et al., 2014). The ligand-receptor complex obtained with molecular docking was immersed in a water box with a 10 Å buffer of water molecules and neutralized by adding explicit Cl^–^ counterions. The region comprised between residues Val306 and Pro332, not resolved in the crystallographic structure of CB_1_, was not modeled, and a chain break was introduced. A two-stage geometry optimization approach was implemented. The first stage minimizes only the positions of solvent molecules and ions, and the second stage is an unrestrained minimization of all the atoms in the simulation cell. The system was then heated by incrementing the temperature from 0 to 300 K under a constant pressure of 1 atm and periodic boundary conditions. Harmonic restraints of 10 kcal mol^-1^ Å^-2^ were applied to the solute, and the Andersen temperature coupling scheme (Andersen, 1980; Andrea et al., 1983) was used to control and equalize the temperature. The time step was kept at 1 fs during the heating stages, allowing potential inhomogeneities to self-adjust. The SHAKE (Miyamoto & Kollman, 1992) algorithm was employed for further equilibration and production with a 2 fs time step. Long-range electrostatic effects were modelled using the particle mesh Ewald method (Darden et al., 1993). A cutoff of 8 Å was applied to Lennard-Jones interactions. The system was equilibrated for 2 ns at constant volume and temperature of 300 K, and production was run as a 100 ns trajectory in the same conditions.

### Statistical analysis

Data from radioligand affinity assays were analyzed using nonlinear regression. Data from autoradiographic assays was evaluated using Kruskal-Wallis test followed by Dunn’s post hoc tests for multiple comparisons. The threshold for statistical significance was set at p < 0.05. Statistics and data were graphically represented using GraphPad Prism 9 (GraphPad Software).

## Results

### Chemical synthesis of the novel compounds

As a follow-up of previously described studies regarding the palladium-catalyzed heterocyclization/oxidative Heck coupling cascade as a synthetic approach to fused heterocycles, we have explored the feasibility of performing the reaction sequence in a one-pot fashion, merging Sonogashira-heterocyclization and oxidative Heck processes in the same step and using the same catalyst, to generate a new series of dihydroindolo[3,2-d]benzazepine-6(5*H*)-ones starting from simple protected *o*-iodoanilines and acyl *o*-alkynylanilines (see figure 1).

**Figure 1.**
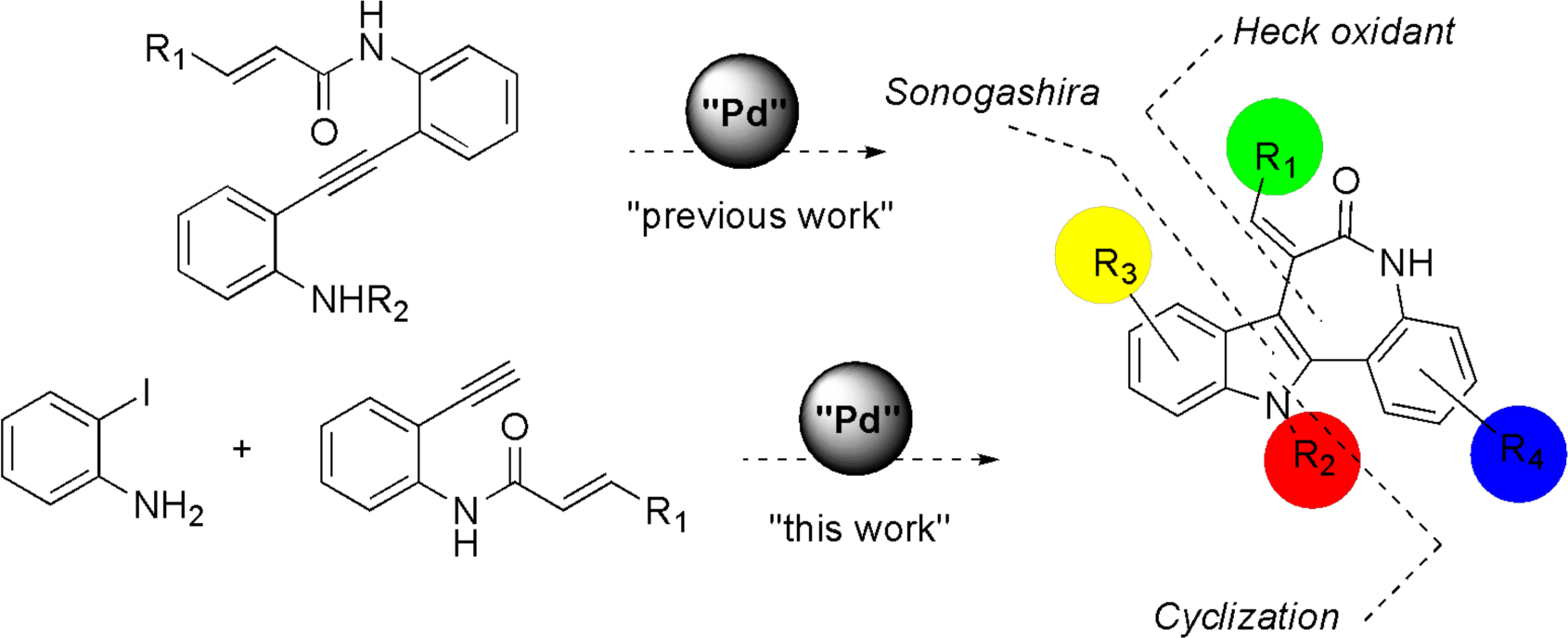
Synthetic approach to the construction of the dihydroindolo[3,2-d]benzazepine-6(5*H*)-one skeleton, including the intra- and intermolecular version.

In the synthetic design we have also considered that the indole derivatives protected as carbamates have better solubility, and they are easier to purify and crystallize than the corresponding free indoles with the same substitution pattern, while the carbamate group does not prevent cyclization through the nitrogen in the nucleopalladation step (Denis et al., 2015).

The synthesis of the precursors included two straightforward reactions shown in figure 2, namely the previously described procedures for 2-haloarylcarbamates **2** (Denis et al., 2015; Sakamoto et al., 1987) and the condensation of the 2-ethynylanilines **3** with the corresponding acid chloride for the acyl derivatives **4**, both proceeding in good yields (Denis et al., 2015; Grigg et al., 2002; Ma et al., 2001).

**Figure 2.**
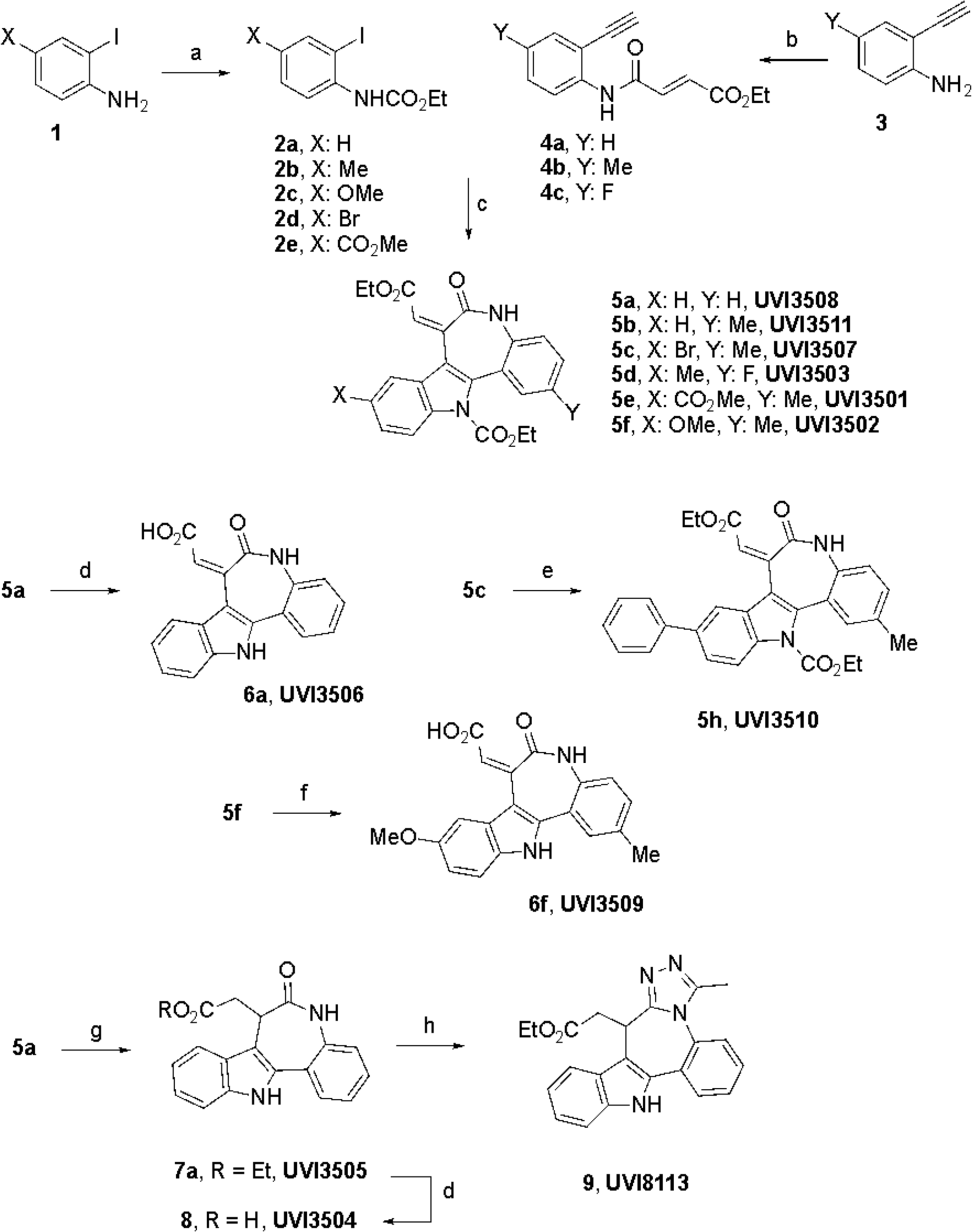
Reagents and reaction conditions: (a) ClCO_2_Et, K_2_CO_3_, acetone, 25 °C (**2a-e**, 74-81%). (b) Ethyl fumaroyl chloride, NaHCO_3_, dioxane, 0 to 25 °C, 23h (**4a-c**, 66-90%). (c) i. PdCl_2_(PPh_3_)_2_ (5 mol%), CuI (20 mol%), PPh_3_ (5 mol%), Et_3_N, DMF, 60 °C, 12h; ii. Air, 100 °C, 15h (**5a-f**, 47-75%). (d) LiOH H_2_O, THF, 50 °C, 12 h (**6a**, 86%; **8**, 79%). (e) PhB(OH)_2_ (1.5 mol equiv), Pd(OAc)_2_ (5 mol%), P(*o*-tol)_3_ (10 mol%), 1,4-dioxane/H_2_O, 80 °C, 12h (**5h**, 50%). (f) TBAF, THF, 80 °C, 21h (**6f**, 71%). (g) i. H_2_, Pd(OH)_2_, EtOAc, 25 °C; ii. 1M TBAF in THF, 80 °C, 21h (**7a**, 73%). (h) Lawesson’s reagent, THF, 60 °C, 3h; ii. H_2_NNH_2_. H_2_O, THF, 20 °C, iii. CH_3_C(OEt)_3_, THF, 80 °C, 1h (**9**, 87%).

Based on the previously reported conditions for the formation of fused indoles in one-pot sequence (Denis et al., 2015), the construction of the target skeleton included: (a) treatment of the corresponding o-iodoaniline 2 (1 equiv) and alkyne 4 (2 equiv) with PdCl_2_(PPh_3_)_2_ (5 mol%) as the catalyst and CuI (20 mol%), Ph_3_P (5 mol%), Et_3_N (2 equiv) as the additives in N,N-dimethylformamide (DMF) under argon atmosphere at 50 °C for 0.5h; (b) opening the flask to air and heating to 50 °C for 17-22h. Indoles 5a-f were obtained in 47-75% yields depending upon the substitution pattern, which can be considered as a very efficient protocol given the increase of structural complexity resulting from three consecutive synthetic steps.

Straightforward synthetic modifications (deprotection and hydrolysis of the ester) allowed to convert the carbamate/ester of **5a** into **6a**, and **5f** into **6f** (see figure 2). In addition, a Pd-catalyzed cross coupling of **5c** with phenylboronic acid in the presence of catalytic quantities of Pd(OAc)_2_ and P(*o*-tol)_3_ in dioxane/H_2_O at 80 °C led to **5h** (see figure 2). Hydrogenation of the conjugated ester upon catalysis of Pd(OH)_2_ in ethyl acetate and deprotection of the carbamate with TBAF afforded **7a** in a combined 73% yield, which was alternatively hydrolyzed to **8**, or converted into the fused triazole **9** in an overall 87% yield upon combined treatment with Lawesson’s reagent at 60 °C, followed by hydrazine hydrate and triethyl orthoacetate in THF at 80 °C (see figure 2).

### Screening of the newly synthesized compounds for cannabinoid receptor affinity

*In vitro* binding assays using a selective radioactively labeled cannabinoid ligand, [^3^H]CP55,940, with varying concentrations of the newly synthesized compounds were performed to determine their affinity to cannabinoid receptor subtypes. Membrane homogenates from Sprague-Dawley rat cortical tissue were used for these assays since the brain tissue was the therapeutic target to develop these compounds. The IC_50_ (nM) values for each compound obtained in these assays are summarized in Supplemental Table 1. Out of the 12 compounds analyzed, two of them, UVI3502 and UVI3510, showed some affinity for cannabinoid receptors. UVI3502 showed a relatively high affinity with two binding sites in the [^3^H]CP55,940 inhibition curve (IC_50_Hi 0.47 ± 1.94 nM and IC_50_Lo 1470 ± 1.80 nM, see figure 3 and supplemental table 1), while UVI3510 showed a single low affinity binding site (IC_50_Lo 1147 ± 1.65 nM, see figure 3 and supplemental table 1). The total inhibition of [^3^H]CP55,940 binding exerted by UVI3502 was about 72%, with approximately 33% corresponding to the high affinity binding site and the remaining 39% corresponding to the low affinity one. Consequently, UVI3502 was chosen as the best candidate compound based on its affinity for cannabinoid receptors.

**Figure 3.**
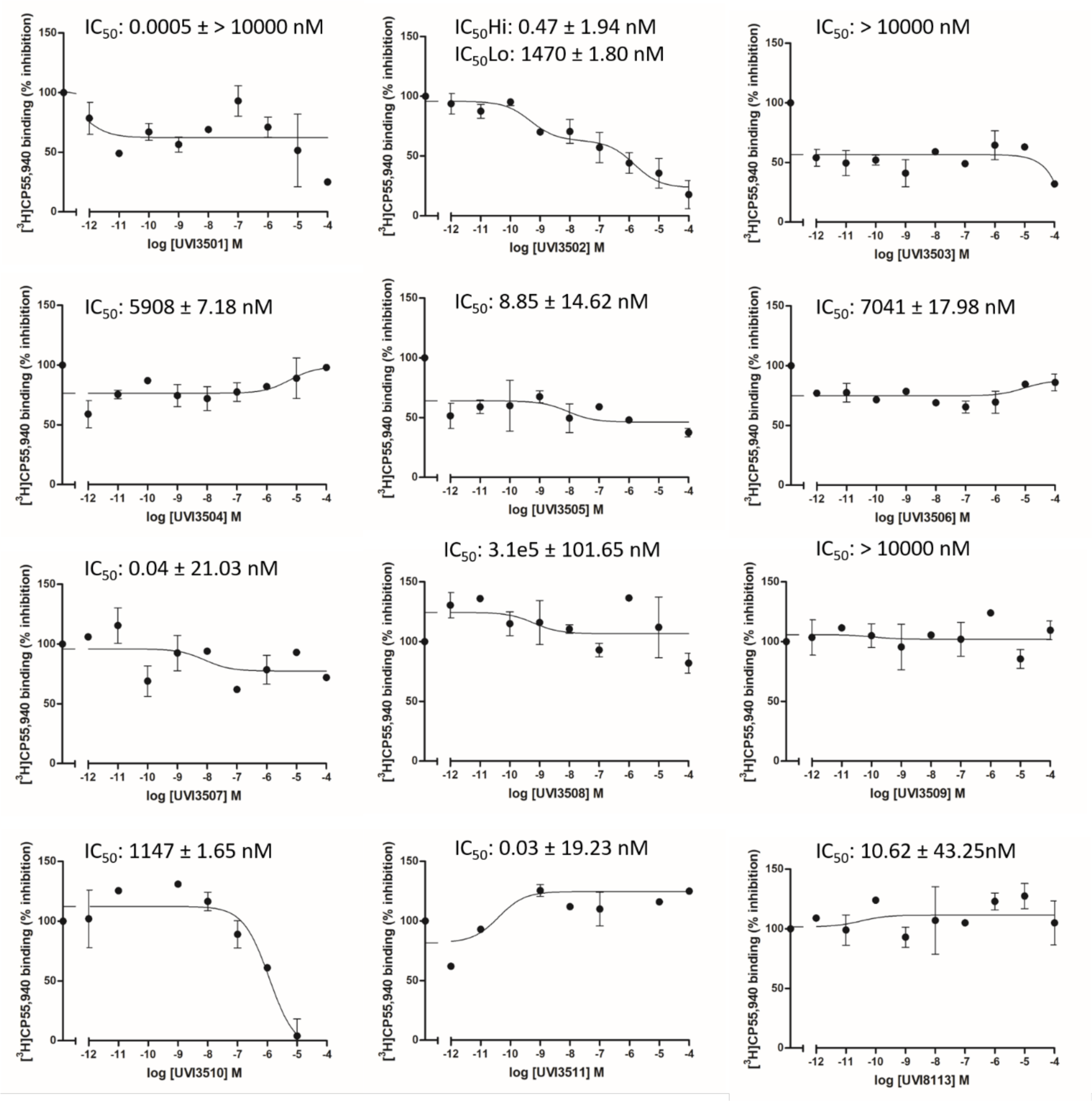
Inhibition curves of [^3^H]CP55,940 *vs.* increasing concentrations ranging from 10^–12^ M to 10^–4^ M of the 12 novel compounds in membrane homogenates from rat brain cortex. Out of the 12 compounds, two of them, UVI3502 and UVI3510, inhibited [^3^H]CP55,940 binding. Note the higher affinity shown by UVI3502 as compared to UVI3510.

### Screening of the affinity of UVI3502 for cannabinoid receptors subtypes

The pharmacological profiling of UVI3502 as a novel cannabinoid ligand was completed by performing inhibition curves of [^3^H]CP55,940 *vs.* a range of concentrations of UVI3502 in membrane homogenates from CHO cells overexpressing CB_1_ and CB_2_ receptors. As a further control, inhibition curves were also performed in rat spleen tissue, due to the high density of CB_2_ receptors and practical absence of CB_1_ receptors in this organ (Pertwee, 1997).

UVI3502 showed affinity for CB_1_ receptors and a single binding site in CB_1_ overexpressing cells (IC50 89.30 ± 1.49 nM, R^2^=0.8717; see figure 4), suggesting that the IC50Lo value observed in the inhibition curve performed in rat brain cortex tissue could correspond to CB_1_ receptors.

**Figure 4.**
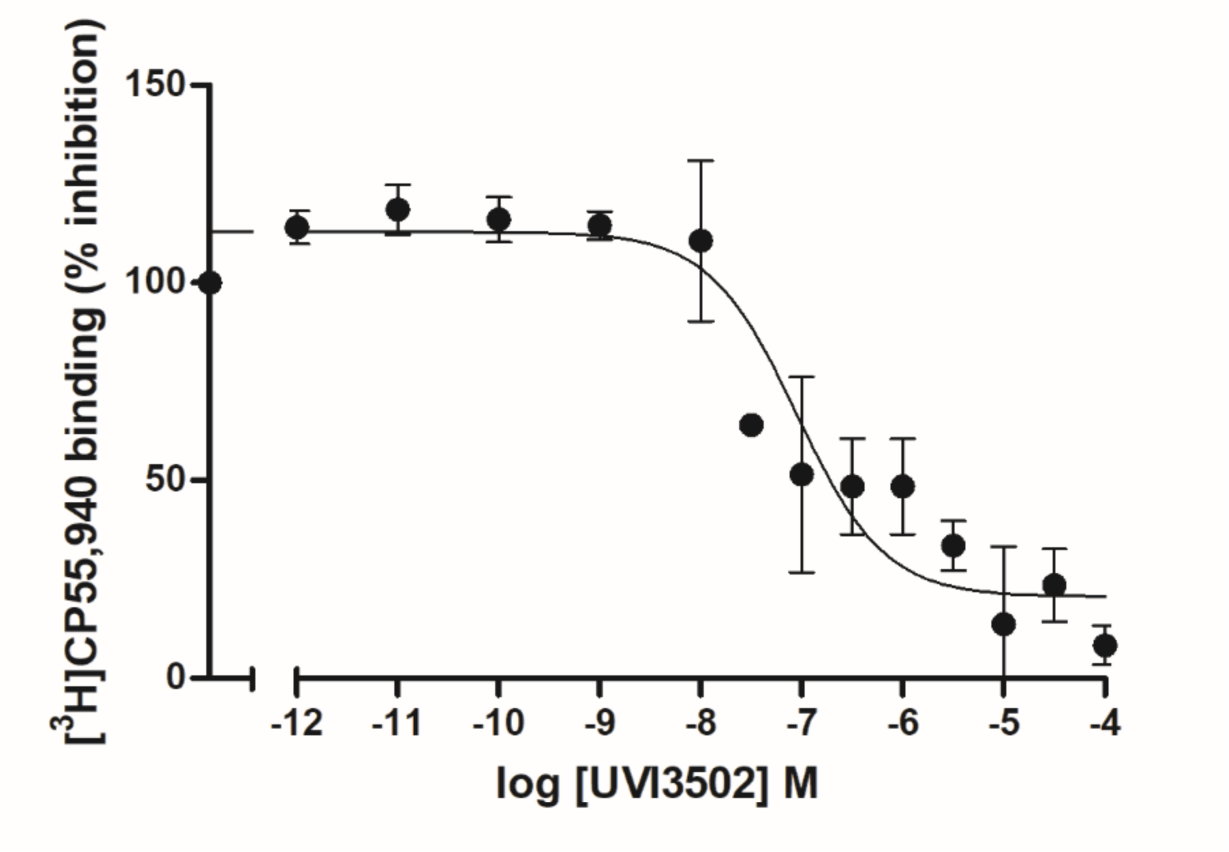
Inhibition curve of [^3^H]CP55,940 *vs.* increasing concentrations ranging from 10^–12^ M to 10^–4^ M of UVI3502 in CB_1_ overexpressing cells. Note that the curve shows a single binding site, indicating that UVI3502 binds CB_1_ receptors (IC_50_ 89.30 ± 1.49 nM, R^2^=0.8717).

In contrast, UVI3502 showed a very low affinity for CB_2_ receptor, only inhibiting [^3^H]CP55,940 binding in CB_2_ overexpressing cells and in spleen membrane homogenates in concentrations at the low micromolar range (CB_2_ overexpressing cells: IC_50_ 8612 ± 13.67 nM, R^2^=0.8854; spleen membrane homogenates: IC_50_ 10230 ± 17.83 nM, R^2^=0.6216; see figure 5). These results suggest that the IC_50Hi_ value observed in the inhibition curve performed in rat brain cortex tissue does not correspond to CB_2_ receptor subtype.

**Figure 5.**
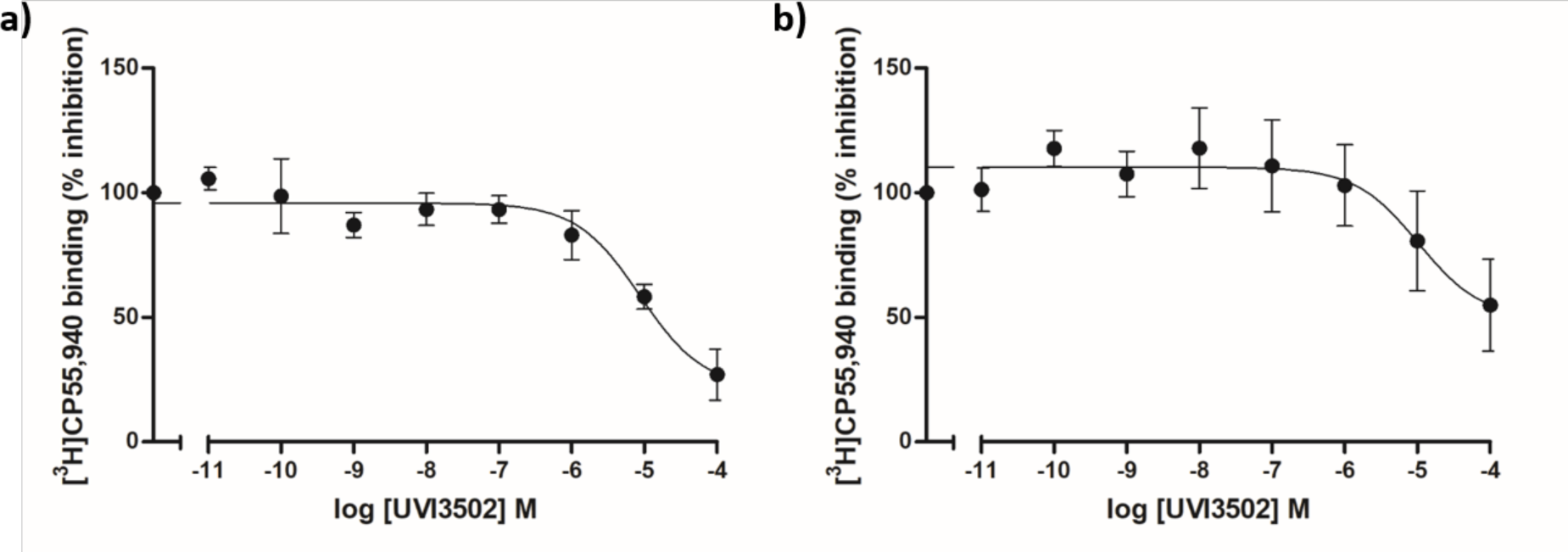
Inhibition curves of [^3^H]CP55,940 *vs.* increasing concentrations ranging from 10^–12^ M to 10^–4^ M of UVI3502 in CB_2_ overexpressing cells and in membrane homogenates from rat spleen. (a) Inhibition curve in CB_2_ overexpressing cells. Note that UVI3502 shows very low affinity for CB_2_ receptor (IC_50_ 8612 ± 13.67 nM, R^2^=0.8854). (b) Inhibition curve in membrane homogenates from rat spleen. Note that UVI3502 shows very low affinity for CB_2_ receptor also in this CB_2_-enriched tissue (IC_50_ 10230 ± 17.83 nM, R^2^=0.6216).

Given that the high affinity binding site observed in the inhibition curve performed in rat brain cortex tissue (see figure 3) does not correspond to CB_1_ or CB_2_ receptor subtypes, we investigated the binding of UVI3502 to GPR55 as the most likely target that could correspond to the high affinity binding site observed, given the described affinity of [^3^H]CP55,940 for this receptor (Ryberg et al., 2007). The unavailability of commercial GPR55 overexpressing membrane homogenates led us to perform inhibition curves using [^3^H]CP55,940 *vs.* increasing concentrations of LPI, the endogenous ligand of GPR55 receptor (Oka et al., 2007), in membrane homogenates from rat brain cortex. LPI inhibited approximately 22% of [^3^H]CP55,940 binding (IC_50_ 0.043 ± 3.39 nM, R^2^=0.5014; see figure 6). The percentage of inhibition and the IC_50_ could correspond to the IC_50Hi_ value obtained by UVI3502 in the [^3^H]CP55,940 inhibition curve, suggesting that the high affinity binding site could correspond to GPR55 receptor. However, a more direct approach, such as an inhibition curve using GPR55 expressing cells with a radiolabeled GPR55 ligand *vs.* UVI3502, is needed to verify this observation.

**Figure 6.**
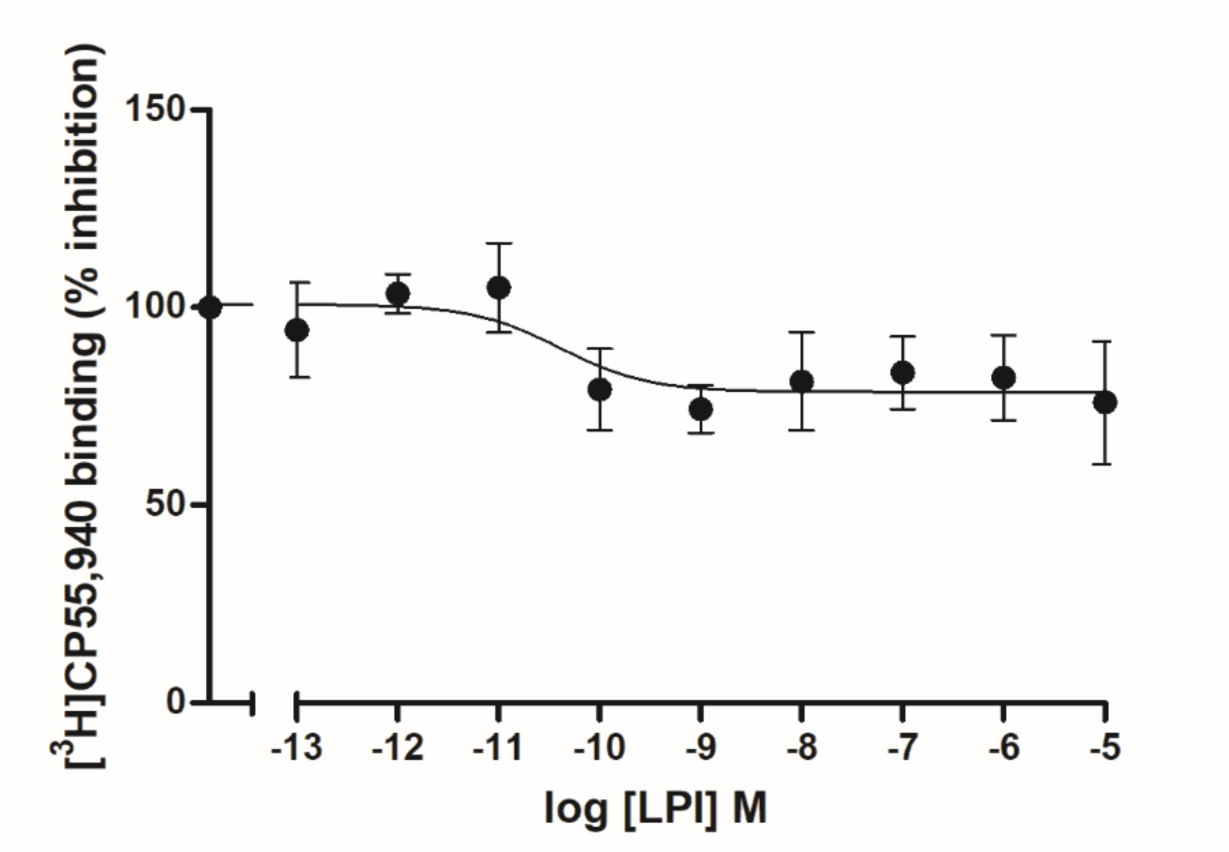
Inhibition curve of [^3^H]CP55,940 *vs.* increasing concentrations of LPI ranging from 10^–13^ M to 10^–5^ M in membrane homogenates from rat brain cortex. Note that the curve shows a binding site, indicating that LPI binds GPR55 receptor (IC_50_ 0.043 ± 3.39 nM, R^2^=0.5014), inhibiting approximately 22% of [^3^H]CP55,940 binding, potentially corresponding to the high affinity binding site observed in the [^3^H]CP55,940 *vs.* UVI3502 inhibition curve in the rat cortex membrane homogenates.

### Characterization of the binding of UVI3502 to CB_1_ receptor with neuroanatomical specificity in the rodent brain

The pharmacological profile of UVI3502 was further characterized in the rodent brain with neuroanatomical specificity by performing [^3^H]CP55,940 autoradiographic assays in brain slices from naïve Sprague-Dawley rats and Swiss mice (see figure 7).

**Figure 7.**
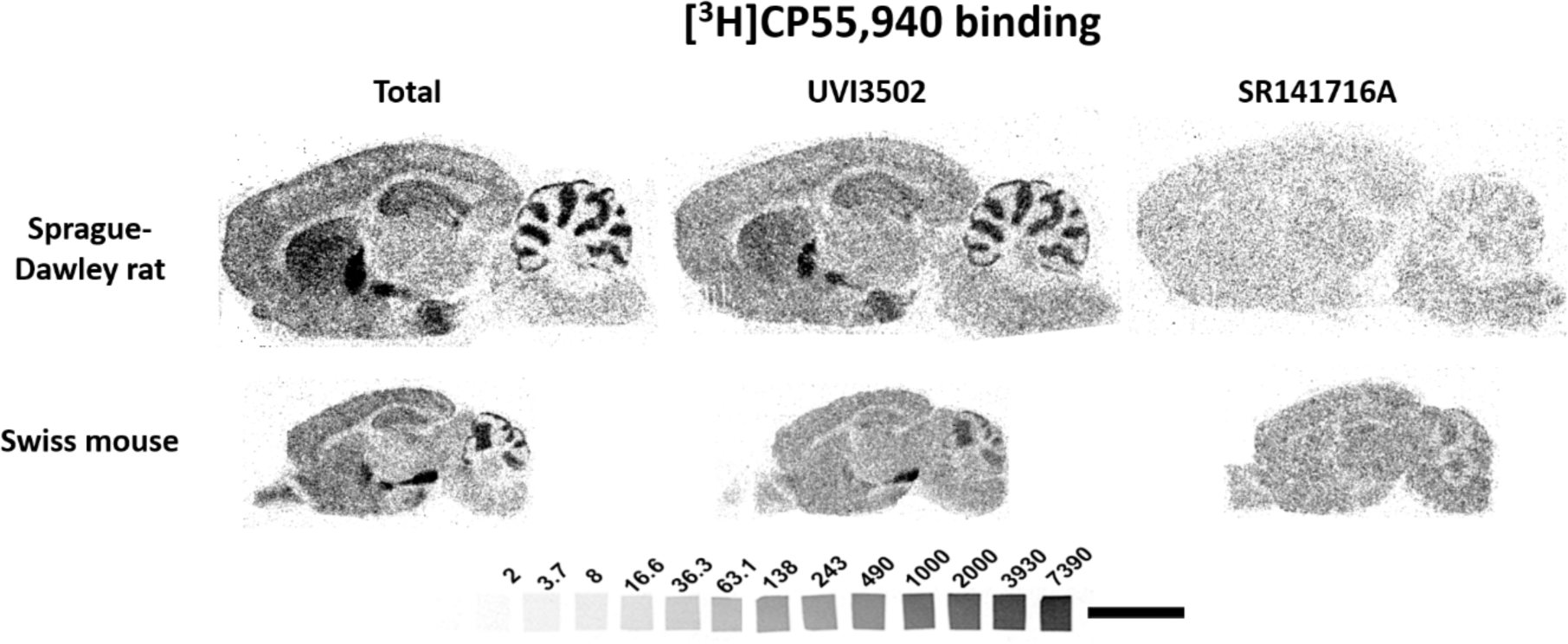
Representative autoradiograms of rats (n = 5) and mice (n = 5) brain sagittal sections showing [^3^H]CP55,940 binding alone, [^3^H]CP55,940 binding in the presence of UVI3502 and [^3^H]CP55,940 binding in the presence of SR141716A. [^3^H] microscales used as standards in Ci/g t.e. Note the partial inhibition of [^3^H]CP55,940 binding by UVI3502 in most of the brain areas expressing CB_1_ receptors. Bar = 0.5 cm.

[^3^H]CP55,940 autoradiography was performed in the presence of both the radioligand and UVI3502, as well as in the presence of the radioligand and SR141716A, a known antagonist/inverse agonist of CB_1_ receptors, in order to have a reference of the amount of [^3^H]CP55,940 binding inhibited by the novel compound. UVI3502 was able to partially inhibit [^3^H]CP55,940 binding in all the areas that were analyzed in both rat and mice brain slices (see figures 8 and 9 and supplemental table 2). These results confirm that UVI3502 binds CB_1_ receptors and indicate that it inhibits a fraction of the [^3^H]CP55,940 binding inhibited by the full antagonist/inverse agonist SR141716A.

**Figure 8.**
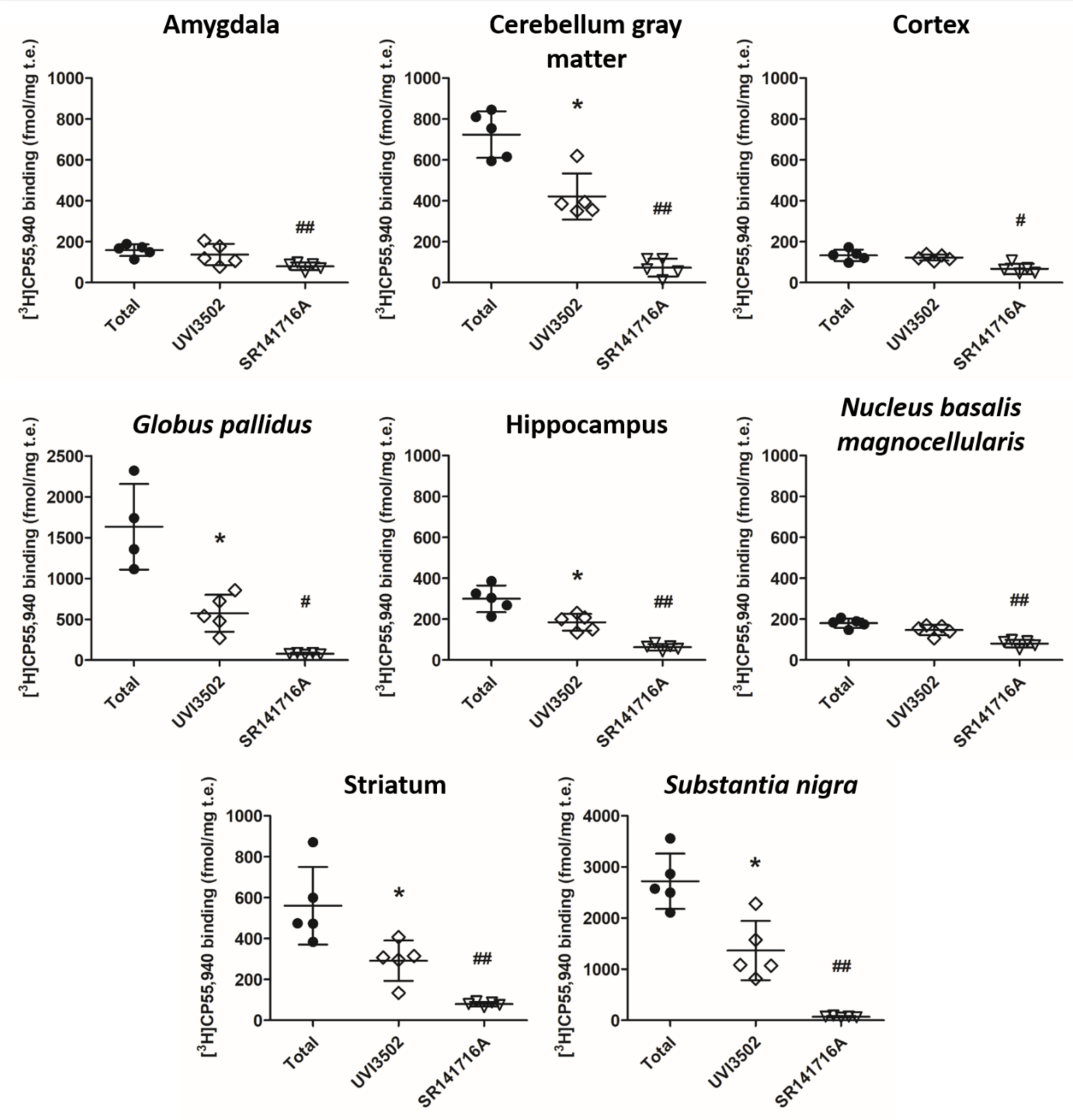
Total [^3^H]CP55,940 binding and [^3^H]CP55,940 binding in the presence of UVI3502 and SR141716A (10 µM) in brain slices from Sprague-Dawley rats (n = 5) in the amygdala, the gray matter of the cerebellum, the cortex, the hippocampus, the *globus pallidus*, the *nucleus basalis magnocellularis* (NBM), the striatum and the *substantia nigra*. SR141716A inhibited [^3^H]CP55,940 binding in all areas, while UVI3502 inhibited a fraction of [^3^H]CP55,940 binding in the gray matter of the cerebellum, the hippocampus, the *globus pallidus*, the striatum and the *substantia nigra* (Kruskal–Wallis test, *post-hoc* test Dunn’s multiple comparison, total *vs.* UVI3502 (*) and total *vs.* SR141716A (^#^), *^#^p < 0.05, ^##^p < 0.01).

**Figure 9.**
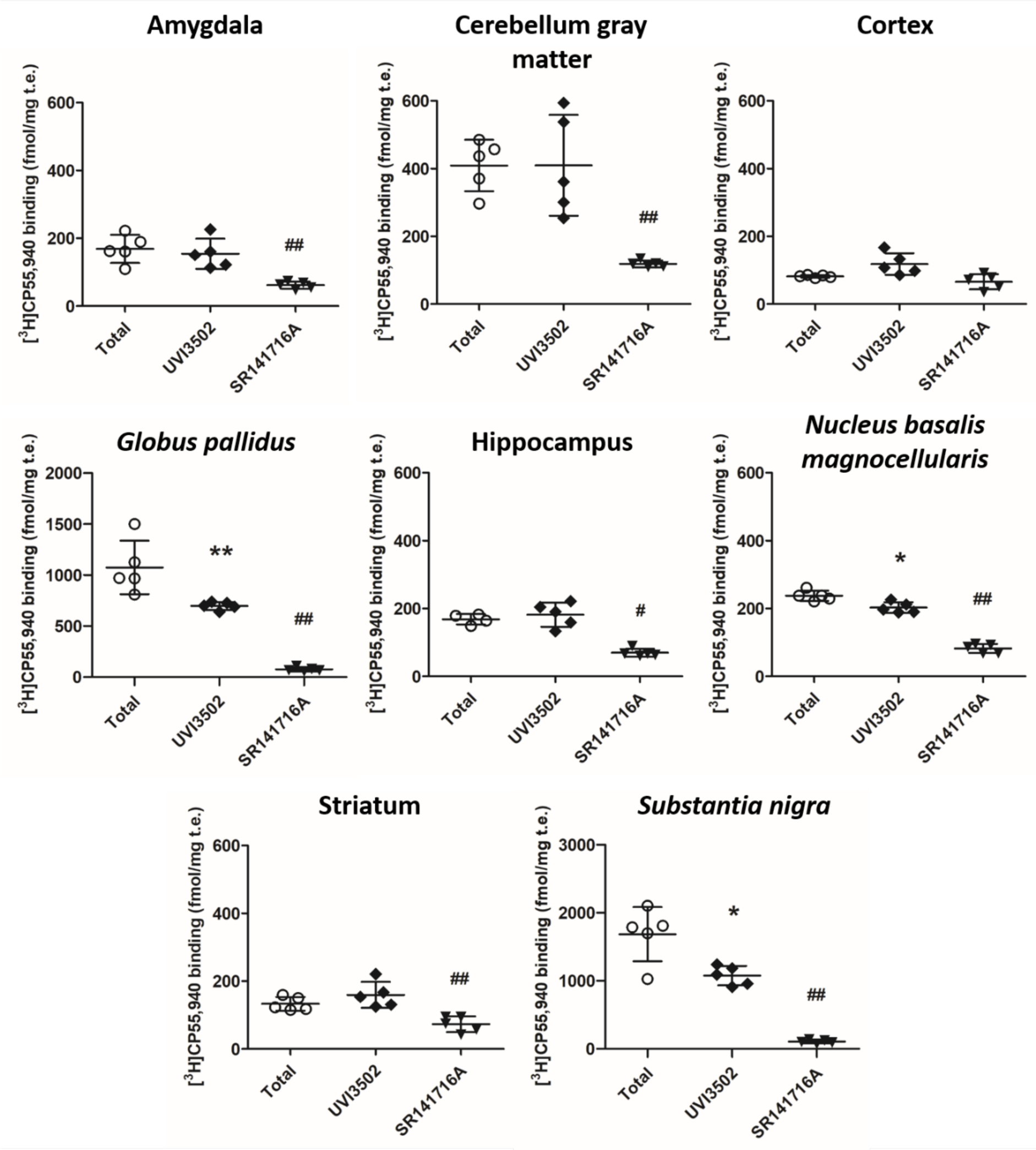
Total [^3^H]CP55,940 binding and [^3^H]CP55,940 binding in the presence of UVI3502 and SR141716A (10 µM) in brain slices from Swiss mice (n = 5) in the amygdala, the gray matter of the cerebellum, the cortex, the hippocampus, the *globus pallidus*, the NBM, the striatum and the *substantia nigra*. SR141716A inhibited [^3^H]CP55,940 binding in all areas, while UVI3502 inhibited a fraction of [^3^H]CP55,940 binding in the *globus pallidus*, the NBM and the *substantia nigra* (Kruskal–Wallis test, *post-hoc* test Dunn’s multiple comparison, total *vs.* UVI3502 (*) and total *vs.* SR141716A (^#^), *^#^p < 0.05, **^##^p < 0.01).

### [^35^S]GTPɣS functional assay to characterize the activity of UVI3502 at CB_1_ receptor

To study the activity of UVI3502 at CB_1_ receptors, a [^35^S]GTPɣS functional assay in membrane homogenates from rat cortical tissue was performed. In this assay, UVI3502 did not stimulate the coupling of CB_1_ receptors to G_i/o_ proteins and in fact decreased baseline coupling levels, starting at nanomolar concentrations. These results suggest that UVI3502 is acting as an antagonist/inverse agonist of CB_1_ receptors (EC_50_ 4.06 ± 3.27 nM, R^2^=0.5992; see figure 10).

**Figure 10.**
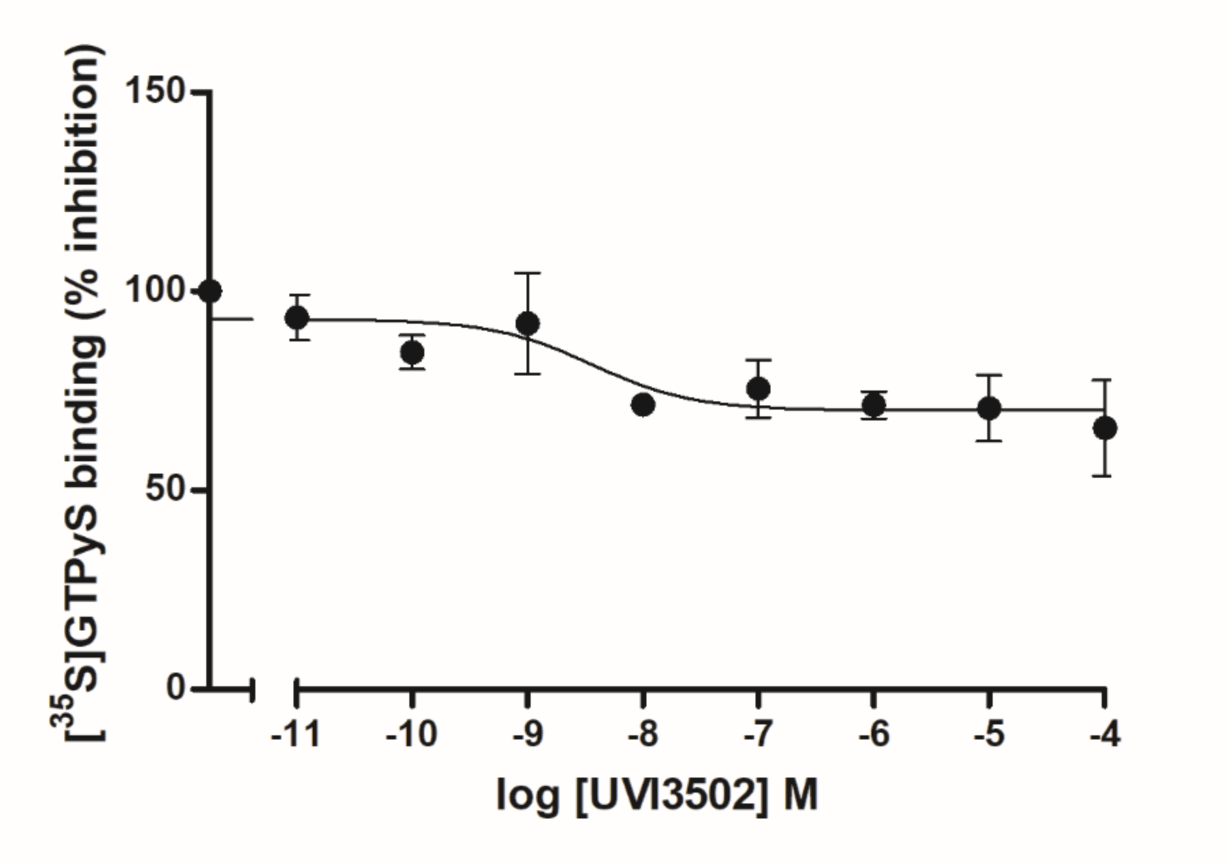
[^35^S]GTPγS binding curve in the presence of UVI3502 in concentrations ranging from 10^–11^ M to 10^–4^ M in membrane homogenates from rat brain cortex. Note that UVI3502 does not stimulate [^35^S]GTPγS binding and in fact decreases basal levels, starting at nanomolar concentrations, indicating it is acting as an inverse agonist of CB_1_ receptors (EC_50_ 4.06 ± 3.27 nM, R^2^=0.5992).

### Characterization of the activity of UVI3502 with neuroanatomical specificity in the rodent brain

The activity of UVI3502 was further analyzed with neuroanatomical specificity by performing [^35^S]GTPɣS autoradiography experiments. The assay was performed by incubating [^35^S]GTPɣS with CP55,940 (10 µM) alone, as a known agonist of CB_1_ receptors, and consecutive slices with UVI3502 (10 µM) alone and in the presence of both CP55,940 (10 µM) and UVI3502 (10 µM) (see figure 11).

**Figure 11.**
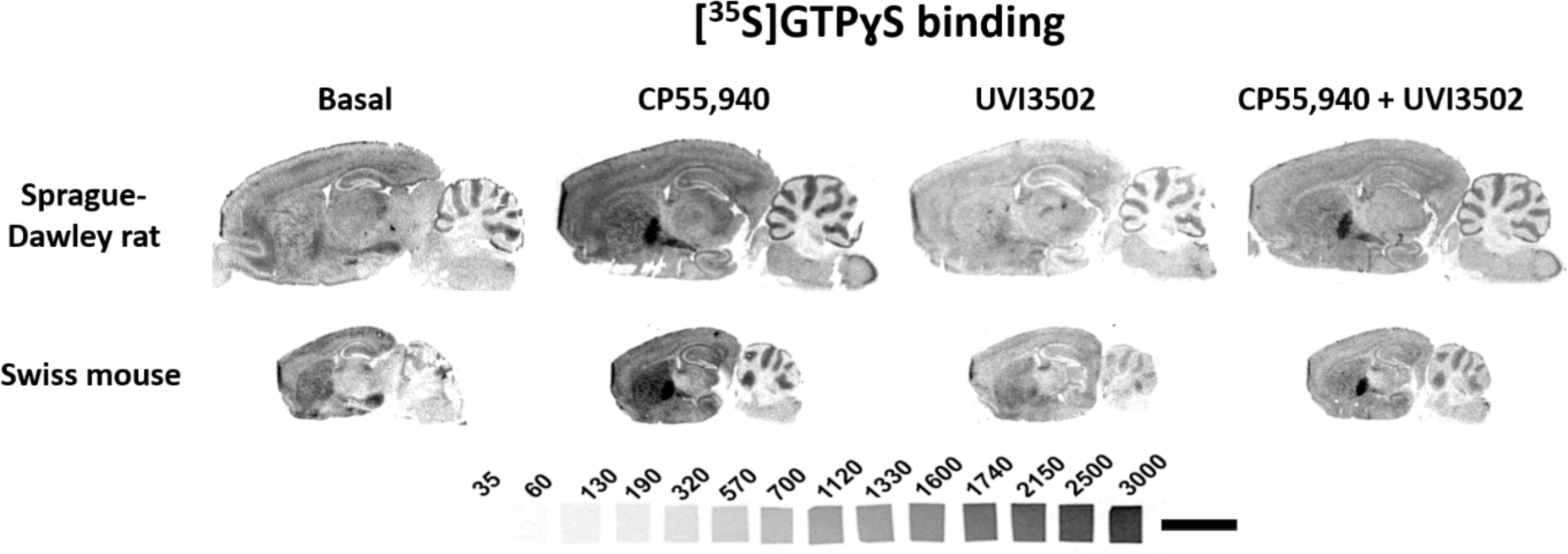
Representative autoradiograms of rats (n = 5) and mice (n = 5) brain sagittal sections that show [^35^S]GTPyS basal binding as well as [^35^S]GTPyS binding in the presence of CP55,940 (10 µM) alone, in the presence of UVI3502 (10 µM) alone and in the presence of both CP55,940 and UVI3502 (both at 10 µM). Note that UVI3502 inhibits the stimulation evoked by CP55,940 in most brain areas and decreases the basal levels of [^35^S]GTPyS binding. [^14^C] microscales used as standards in Ci/g t.e. Bar = 0.5 cm.

Results from Sprague-Dawley rat brain slices confirm that UVI3502 acts as an antagonis/inverse agonist in most of the analyzed areas, as [^35^S]GTPɣS binding in the presence of UVI3502 is lower than baseline levels of [^35^S]GTPɣS binding in the amygdala, the cortex, the hippocampus, the NBM, the striatum and the gray matter of the cerebellum (see figure 12 and supplemental table 3). When UVI3502 was incubated in the presence of CP55,940, the stimulation evoked by the agonist was suppressed in all the analyzed areas (see figure 12 and supplemental table 3).

Regarding the data obtained from mouse brain slices, UVI3502 also acts as an antagonist/inverse agonist in most of the brain areas, decreasing baseline levels of [^35^S]GTPɣS binding (see figure 13 and supplemental table 3). In the presence of UVI3502, the stimulation evoked by CP55,940 was suppressed in the analyzed areas (see figure 13 and supplemental table 3). These results further confirm that UVI3502 acts as a partial antagonist/inverse agonist of CB_1_ receptors in rodent brain tissue.

**Figure 12.**
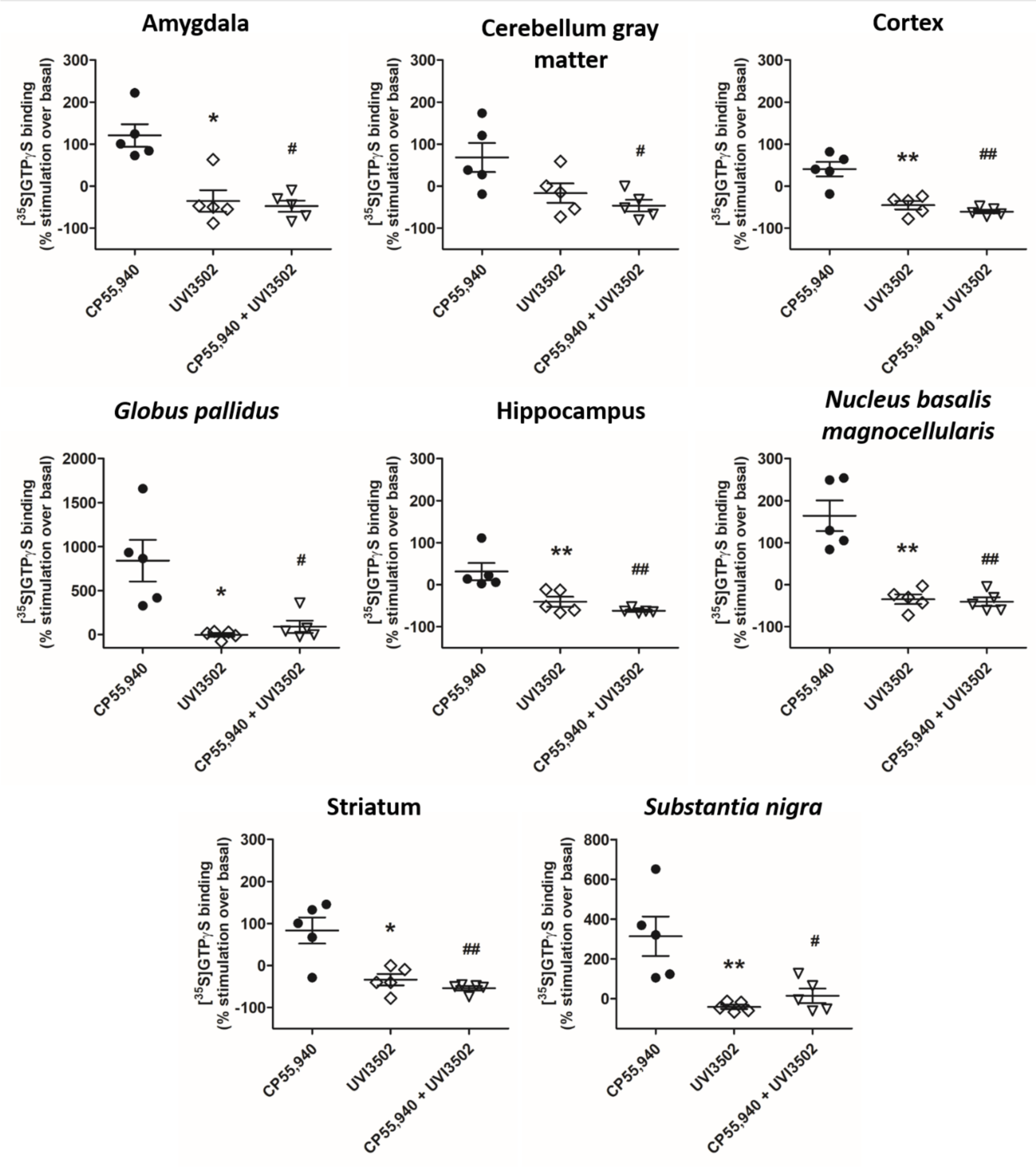
[^35^S]GTPγS binding stimulated by CP55,940 (10 µM), as a reference, and in the presence of UVI3502 (10 µM) and CP55,940 + UVI3502 (both at 10 µM), in brain slices from Sprague-Dawley rats (n = 5) in the amygdala, the gray matter of the cerebellum, the cortex, the hippocampus, the *globus pallidus*, the NBM, the striatum and the *substantia nigra*. UVI3502 decreased basal binding in the amygdala, the cortex, the hippocampus, the *globus pallidus*, the NBM, the striatum and the *substantia nigra*. UVI3502 prevented the stimulation evoked by CP55,940 in all the areas that were analyzed (Kruskal–Wallis test, *post-hoc* test Dunn’s multiple comparison, CP55,940 *vs.* UVI3502 (*) and CP55,940 *vs.* CP55,940 + UVI3502 (^#^), *^#^p < 0.05, **^##^p < 0.01).

**Figure 13.**
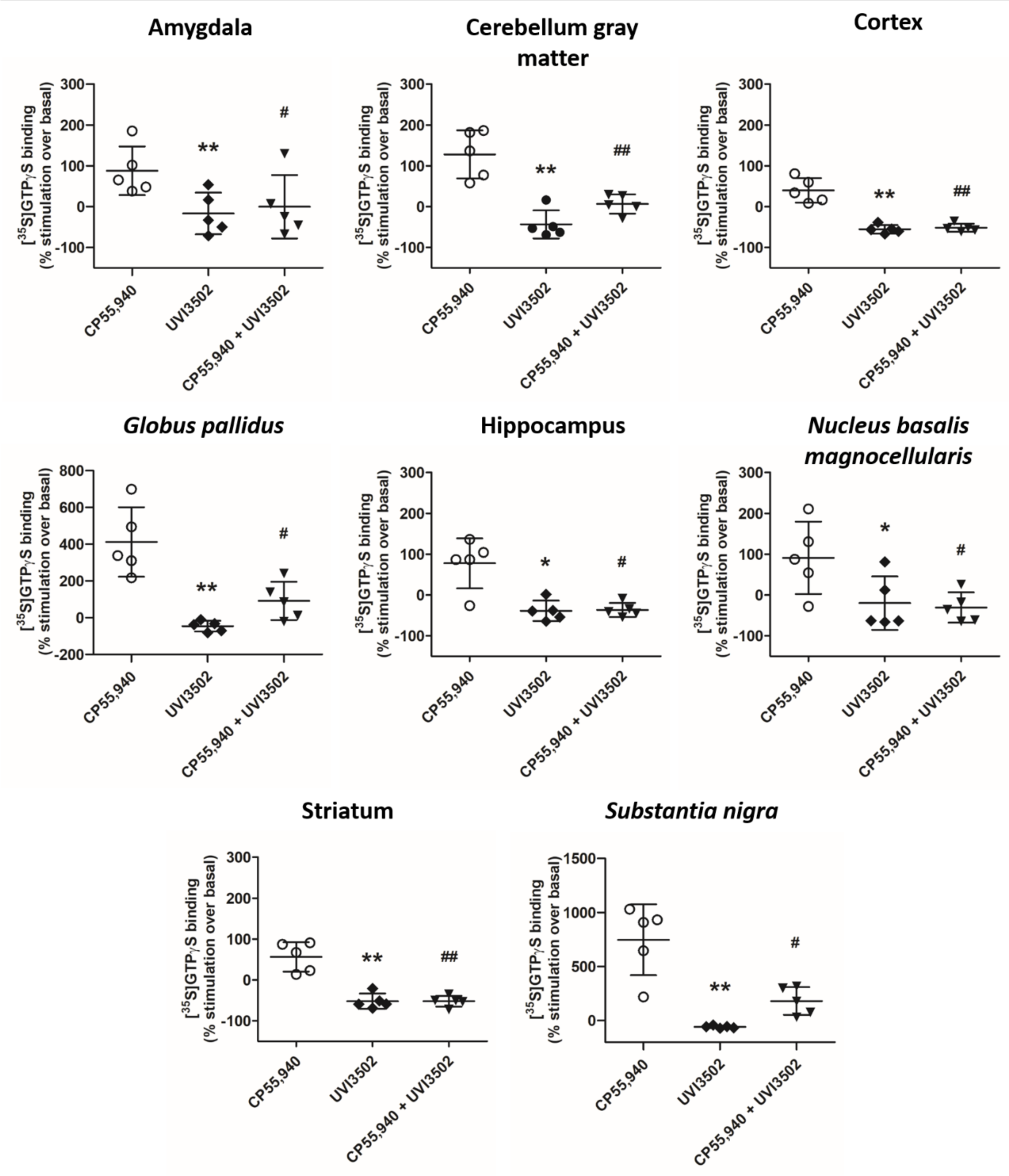
[^35^S]GTPγS binding stimulated by CP55,940 (10 µM), as a reference, and in the presence of UVI3502 (10 µM) and CP55,940 + UVI3502 (10 µM), in brain slices from Swiss mice (n = 5) in the amygdala, the gray matter of the cerebellum, the cortex, the hippocampus, the *globus pallidus*, the NBM, the striatum and the *substantia nigra*. UVI3502 decreased basal binding in the amygdala, the gray matter of the cerebellum, the cortex, the hippocampus, the *globus pallidus*, the NBM, the striatum and the *substantia nigra*. UVI3502 prevented the stimulation evoked by CP55,940 in all the areas analyzed (Kruskal–Wallis test, *post-hoc* test Dunn’s multiple comparison, CP55,940 *vs.* UVI3502 (*) and CP55,940 *vs.* CP55,940 + UVI3502 (^#^), *^#^p < 0.05, **^##^p < 0.01).

### Modeling of UVI3502 binding to CB_1_

Binding of UVI3502 to human CB_1_ receptor was modeled through a combination of molecular docking and classical molecular dynamics. UVI3502 was docked on a three-dimensional model of CB_1_ generated from its crystallographic structure in complex with the known antagonist AM-6538 (see figure 14a) (Hua et al., 2016b). Figure 14b shows the best scoring docking pose (score = 35.5); like AM-6538, UVI3502 is deeply buried in the binding pocket of CB_1_, roughly occupying the same region, and is engaged in hydrophobic contacts with Phe170, Phe174, Phe268, Trp356, and Phe379. The binding pose is driven by a tight shape complementarity (see figure 14c); the main differences observed between the two ligands are a reduced occupation of the long channel (lined by Trp279 and Met363) for UVI3502 owing to the size of the carbamate moiety, and increased interactions with the upper part of the pocket (Phe268 and Phe 379) through the tricyclic core and methoxy group. The two ligands fit the gap and side pocket regions to a similar extent. A 100 ns classical molecular dynamics simulation of the UVI3502:CB_1_ complex was performed using the docking pose as the starting geometry to evaluate the persistence of key binding contacts. Figure 14d shows an overlay of 5 frames sampled with an even stride from the molecular dynamics simulation. Although some flexibility is observed, the positioning and orientation of UVI3502 in the binding pocket remains constant, as well as the packing of the surrounding hydrophobic residues, corroborating the docking pose as a plausible representation of the binding interaction.

**Figure 14.**
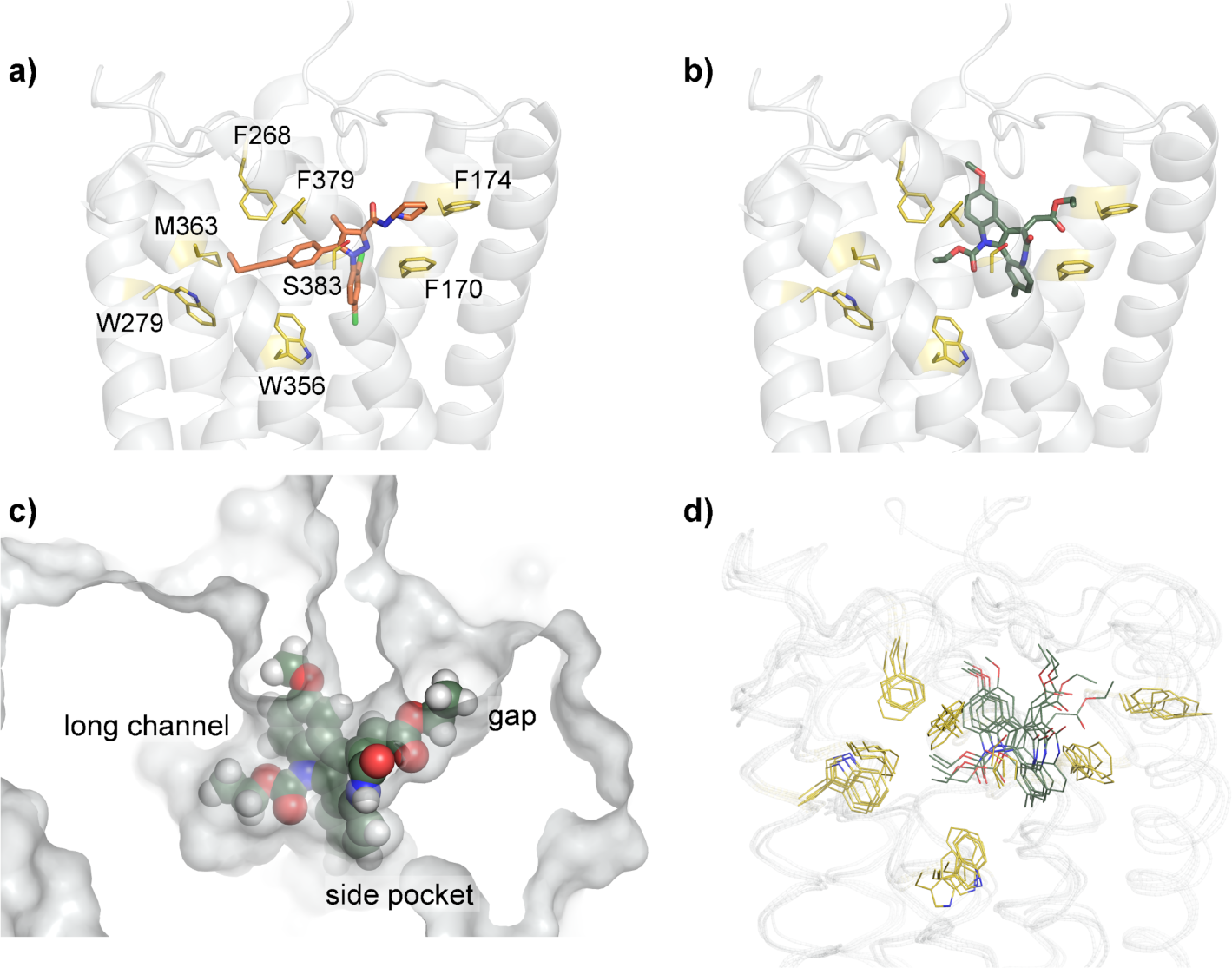
(a) Crystal structure of the human cannabinoid receptor CB_1_ in complex with inverse agonist AM6538 (in pink sticks; terminal nitrate group is not shown due to the absence of electron density), PDB 5TGZ. (b) Docking pose of UVI3502 (in green sticks) on human cannabinoid receptor CB_1_ (receptor structure taken from PDB 5TGZ). (c) Space-filling view of the same docking pose with indication of the main regions involved in UVI3502 binding. (d) Overlay of 5 snapshots sampled with an even stride of 25 ns from a molecular dynamics simulation of the UVI3502:CB_1_ complex. Key residues for inverse agonist binding are shown as yellow sticks.

## Discussion

In light of the therapeutic potential of the eCB system, in particular regarding various neurological disorders, it is essential to develop new compounds that exhibit affinity for cannabinoid receptors. In this study, a series of novel compounds were synthesized and pharmacologically characterized, *in vitro* and *in silico*.

For the synthesis of the compounds, we have increased further the efficiency of the intramolecular oxidative Heck cascade reaction (Denis et al., 2015), by incorporating a Sonogashira cross-coupling prior to the heterocyclization–Heck, as described for other heterocycles, which led to the formation and *N*-cyclization of *o*-alkynylaniline intermediates starting from appropriately protected *ortho*-iodoanilines and terminal alkynylanilines. The sequence of chemical transformations took place in the same reaction flask while additional reagents and catalysts were added at different time intervals (Calder et al., 2014), another example of the ‘one-pot’ multi-component reaction (MCR) (Weber, 2002). This combination of consecutive Sonogashira, nucleopalladation and oxidative Heck couplings conveniently allowed the preparation of a new series of 7,12-dihydroindolo[3,2-d]benzazepine-6(5*H*)-ones. The skeleton of the indolobenzazepinones was further modified by incorporation of additional substituents or by its conversion into the fused [1,2,4]triazoloazepines in an efficient manner.

Following the synthesis, we evaluated the pharmacodynamic parameters of the compounds using radioligand affinity assays with [^3^H]CP55,940 performed in membrane homogenates purified from rat cortical tissue, which naturally contain CB_1_, CB_2_ and GPR55, three cannabinoid or cannabinoid-like receptors recognized by [^3^H]CP55,940 (Ryberg et al., 2007; Schatz et al., 1997). Out of the 12 compounds analyzed, one of them, UVI3502, showed significant affinity for the binding sites of [^3^H]CP55,940 in rat brain cortex. The inhibition curves showed a high affinity (IC_50_ values in the nanomolar range) and a low affinity (IC_50_ value in the micromolar range) binding site, which, given the tissue used for the assay and the pharmacological profile of [^3^H]CP55,940, are likely to correspond to CB_1_, CB_2_ and/or GPR55 receptors. The affinity of UVI3502 was confirmed for CB_1_ and discarded for CB_2_ following inhibition curves performed in cells overexpressing human CB_1_ or CB_2_ receptors. Assays performed in membrane homogenates derived from rat spleen tissue, which has a high density of CB_2_ receptors and practical absence of CB_1_ receptors (Pertwee, 1997), further confirmed the cannabinoid receptor subtype selectivity of UVI3502. The affinity of UVI3502 shows a single binding site in cells overexpressing CB_1_ receptors, suggesting that CB_1_ receptor might correspond to the low affinity binding site observed in the inhibition curve performed in rat cortical tissue, while the high affinity binding site might correspond to a third, non-CB_1_/CB_2_ cannabinoid-like receptor, such as GPR18 or GPR55 (Ramírez-Orozco et al., 2019). However, results obtained from membrane homogenates from rat spleen tissue, where GPR18 is abundantly expressed (Gantz et al., 1997), discard this receptor as a likely target of UVI3502. Given these results, GPR55 receptor is the most likely non-CB_1_ target for UVI3502. To confirm this, and given the unavailability of commercial GPR55 overexpressing cell membrane homogenates, an alternative approach was used, which consisted in conducting inhibition curves of [^3^H]CP55,940 binding in the presence of LPI, a proposed endogenous ligand of the GPR55 receptor (Oka et al., 2007), in membrane homogenates from rat brain cortex. Results showed a single high affinity binding site (IC_50_ value in the nanomolar range), inhibiting approximately 22% of the binding. Both the IC_50_ value and the percentage of inhibition approximately align with the IC_50Hi_ value obtained in the [^3^H]CP55,940 *vs*. UVI3502 inhibition curve in membrane homogenates from rat cortical tissue, suggesting that the high-affinity binding site could be associated with the GPR55 receptor. However, this is an indirect approach which cannot confirm this hypothesis, and a more direct method such as conducting an inhibition curve employing GPR55-expressing cells with a radiolabeled GPR55 ligand (currently unavailable) *vs*. UVI3502 is required. If binding to GPR55 was confirmed, UVI3502 would be part of a rare class of ligands with affinity for CB_1_ and GPR55 receptors, but not CB_2_. To the best of our knowledge, only AM251 and AM281 exhibit this kind of pharmacological profile (Kapur et al., 2009), but both of these drugs exhibit either higher affinity for CB_1_ or similar affinity for both receptors (Harding et al., 2023), while the opposite is true for UVI3502, potentially making its pharmacological profile potentially unique among the various ligands targeting cannabinoid receptors.

Autoradiographic assays using [^3^H]CP55,940 were then conducted on brain slices from naïve Sprague-Dawley rats and Swiss mice to assess the pharmacological profile of UVI3502 in brain regions associated with learning and memory. UVI3502 partially inhibited [^3^H]CP55,940 binding in the examined brain areas of both rats and mice. These results confirm that UVI3502 binds CB_1_ receptors and suggest that it is acting as a partial compound, inhibiting only a fraction of the total [^3^H]CP55,940 binding.

Then, to determine the activity of UVI3502 in CB_1_ receptors, a [^35^S]GTPɣS assay was performed in membrane homogenates from rat brain cortex. UVI3502 did not stimulate the coupling to G_i/o_ proteins and in fact reduced the baseline coupling levels starting at nanomolar concentrations, indicating that UVI3502 behaves as an antagonist/inverse agonist of CB_1_ receptors.

In a subsequent functional autoradiographic assay to determine the activity of UVI3502 in key brain areas from the rodent brain, UVI3502 also exhibited the characteristics of an antagonist/inverse agonist across most of the examined regions. This was evident from the lower [^35^S]GTPɣS binding in the presence of UVI3502 compared to baseline levels of [^35^S]GTPɣS binding in areas such as the amygdala, the cortex, the hippocampus, the NBM, the striatum and the cerebellar gray matter. The interpretation of basal [^35^S]GTPɣS binding in the absence of any drug, and thus of the concept of “inverse agonist”, in *in vitro* assays remains a subject of controversy. While numerous reports suggest that different receptors can be constitutively active in the absence of any ligand (Damian et al., 2012; Muneta-Arrate et al., 2020; Sirohi & Walker, 2015), other reports suggest roles of endogenous ligands in so-called basal activity, such as the formation of adenosine during incubation (Laitinen, 1999), or the presence of endogenous lysophosphatidic acid (LPA) activating LPA_1_ receptors (González de San Román et al., 2015). The pharmacological profile of UVI3502 was further confirmed when incubated along with potent cannabinoid receptor agonist CP55,940, since UVI3502 inhibited the stimulation induced by CP55,940 in all examined regions. This further suggests the role of UVI3502 as an antagonist/inverse agonist of CB_1_ receptors (Berg & Clarke, 2018), given that this compound binds this receptor subtype, which is one of the most expressed GPCRs in the cortex (Herkenham et al., 1991). The activity of UVI3502 as an antagonist/inverse agonist is noteworthy, given that it shares some structural resemblance to the aminoalkylindole WIN55,212-2, a potent cannabinoid receptor agonist (Compton et al., 1992). The *in silico* characterization of UVI3502 binding to CB_1_ receptor through molecular docking and molecular dynamics can tentatively explain this activity in terms of the shape complementarity with the binding pocket in the active and inactive states of the receptor (see figure 15). Indeed, UVI3502 is predicted to bind in a similar arrangement to previously reported inverse agonist AM-6538 (Hua et al., 2016b), extending into the side pocket through its vertical axis and joining the long channel and gap region along its longitudinal axis. The latter is a characteristic feature shared by other known inverse agonists/antagonists (e.g. AM-251, rimonabant) (Bialuk & Winnicka, 2011) and is key for optimal interactions with the receptor in the inactive state (see figure 15), in which it matches the roughly T-shaped binding cavity. For UVI3502, this is enabled by the quite planar and rigid framework extending from the carbamate to the exocyclic double bond. Conversely, known agonists, such AM-11542 (Hua et al., 2017) and WIN55,212-2 bind with an angular shape often emerging from flexible alkyl and aromatic side chains (see figure 15), which in turn provides optimal contacts with a matching roughly V-shaped cavity in the less longitudinally extended active state. Owing to its rigid framework, we hypothesize that UVI3502 cannot adopt such angular shape, providing a molecular basis for its activity and providing insights into the structural nuances of CB_1_ activation.

**Figure 15.**
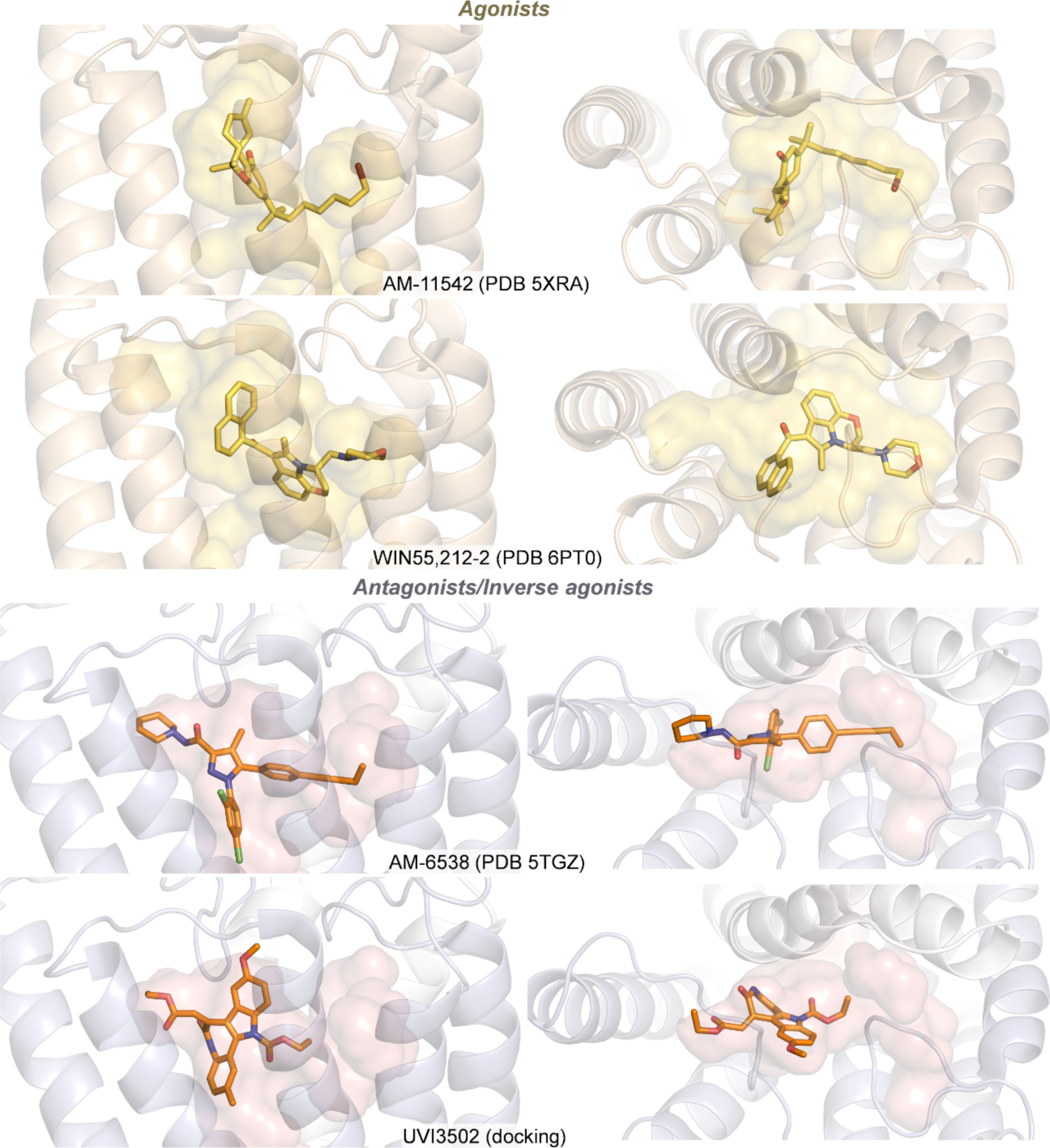
Side view (left) and top view (right) of agonists and inverse agonists/antagonists (AM-11542, WIN55,212-2, AM-6538) bound to human CB_1_ receptors from crystallographic structures and docking pose for UVI3502. The binding cavity is shown as transparent surfaces. Agonists are shown in yellow (top) and antagonists/inverse agonists in orange (bottom).

In summary, we determined the antagonist/inverse agonist properties of the novel compound UVI3502 to CB_1_ receptors, using both *in vitro* and *in silico* approaches. UVI3502 behaves as a partial antagonist/inverse agonist and shows moderate affinity for CB_1_ receptor. Importantly, UVI3502 shows negligible affinity for CB_2_, making it receptor-subtype specific. Moreover, UVI3502 shows high affinity for an additional target yet to be fully determined, which may potentially be the GPR55 receptor. Via functional autoradiography, we determined that UVI3502 blocks the coupling of CB_1_ to G_i/o_ proteins in key brain areas controlling learning and memory, opening the door for the therapeutic use of this compound as a novel inhibitor of cannabinoid receptors.

## Data Availability Statement

The data that support the findings of this study are available on request from the corresponding author. The data are not publicly available due to privacy or ethical restrictions.

## Authorship Contributions

Participated in research design: Bengoetxea de Tena I, Rodríguez De Lera A, Rodríguez-Puertas R.

Conducted experiments: Bengoetxea de Tena I, Pereira-Castelo G, Martínez-Gardeazabal J, Moreno-Rodríguez M, Manuel I, Martínez C, Vaz B, González-Ricarte J, Álvarez R, Torres-Mozas A, Peccati F.

Performed data analysis: Bengoetxea de Tena I, Jiménez-Osés G.

Wrote or contributed to the writing of the manuscript: Bengoetxea de Tena I, Jiménez-Osés G, Rodríguez De Lera A, Rodríguez-Puertas R.

## List of nonstandard abbreviations

Δ9-THC: Δ9-tetrahidrocannabinol
[^35^S]GTPɣS: [^35^S]Guanosine 5’-O-3-thiotriphosphate
AD: Alzheimer’s disease
BSA: Bovine serum albumin
CB_1_: Cannabinoid receptor 1
CB_2_: Cannabinoid receptor 2
CNS: Central nervous system
DS: Down’s syndrome
DTT: DL-dithiothreitol
eCB: Endocannabinoid
GPCR: G-protein coupled receptor
GTPɣS: Guanosine 5’-O-3-thiotriphosphate
HD: Huntington’s disease
LPA: Lysophosphatidic acid
LPI: Lysophosphatidylinositol
MCR: Multi-component reaction
NBM: *Nucleus basalis magnocellularis*
PD: Parkinson’s disease

## Footnotes

a) This work was supported by grants from the Basque Government IT975-16 and IT1454-22 to the “Neurochemistry and Neurodegeneration” consolidated research group; by Instituto de Salud Carlos III through the project PI20/00153 (co-funded by European Regional Development Fund “A way to make Europe”); by MCIN/AEI/10.13039/501100011033 (PID2021-125946OB-I00, RYC2022-036457-I) and by BIOEF through the project BIO22/ALZ/010 funded by Eitb Maratoia. I.B.d.T. is the recipient of an Investigo fellowship funded by the European Union-Next Generation EU. G.P-C. is the recipient of a University of the Basque Country predoctoral fellowship. J.M-G. is the recipient of a Margarita Salas fellowship funded by the European Union-Next Generation EU.

b) Doctoral thesis: Bengoetxea de Tena, I. (2024). *Cannabinoid receptors in animal models of cognitive impairment.* University of the Basque Country (UPV-EHU). Meeting abstract: Bengoetxea de Tena, I. (2021). A new synthetic antagonist for CB_1_ receptor shows affinity for two binding sites of CP55,940. 21a Reunión Anual de la Sociedad Española de Investigación sobre Cannabinoides. Málaga, España: Universidad de Málaga.

c) Rafael Rodríguez-Puertas Department of Pharmacology, Faculty of Medicine and Nursing, University of the Basque Country (UPV/EHU), Leioa, Spain

## Notes

### Competing Interest Statement

The authors have declared no competing interest.

