## Supplemental data for "Synthesis and pharmacological characterization of UVI3502, a novel cannabinoid receptor 1 (CB_1_) antagonist/inverse agonist"

**General information for the synthesis of the novel compounds**

THF was dried using a Puresolv solvent purification system. Commercial anhydrous 1,4-dioxane (99.8%) and DMF (≥99.8%) were kept over 4 Å MS. All other reagents were commercial compounds of the highest purity available. For reactions that require heating we used the Heat-On block system of Radleys. New compounds were fully characterized by their $^{1}H$ and $^{13}C$ NMR, IR, and HRMS spectral properties. Unless otherwise indicated, routine NMR spectra were obtained at 25 °C on a Bruker ARX-400 spectrometer (400.16 MHz for $^{1}H$ and 100.62 MHz for $^{13}C$) using $CDCl_3$, $CD_3OD$, and DMSO-$d_6$ as solvents and internal reference ($CDCl_3$ δ 7.26 for $^{1}H$ and δ 77.0 for $^{13}C$, $CD_3OD$ δ 3.31 for $^{1}H$ and δ 49.0 for $^{13}C$, DMSO-$d_6$ δ 2.50 for $^{1}H$ and δ 39.5 for $^{13}C$). Chemical shifts (δ) are reported in ppm and coupling constants (*J*) are given in hertz (Hz). The proton spectra are reported as follows: multiplicity, coupling constant *J*, number of protons. The DEPT sequence was routinely used for $^{13}C$ multiplicity assignment. Multiplicity in the $^{13}C$ NMR spectral data refers to the attached hydrogens. Additionally, a combination of COSY, HSQC, and HMBC NMR experiments were used for structural assignments. Infrared spectra (IR) data were obtained from a thin film deposited onto a NaCl glass and were measured on a Jasco FT/IR 4100 in the interval between 4000 and 600 $cm^{-1}$ with a 4 $cm^{-1}$ resolution; data include only characteristic absorptions. Electrospray ionization (ESI) mass spectra were obtained on a micrOTOF focus mass spectrometer (Bruker Daltonics) using an ApolloII (ESI) source with a voltage of 4500 V applied to the capillary. Electron impact (EI) mass spectra were obtained on a Hewlett-Packard HP59970 instrument operating at 70 eV. For UPLC-QTOF, chromatographic separation was done with an Acquity UPLC BEH C18 1.7 µm, 50 mm × 2.1 mm column, and $H_2O/HCO_2H$ (99.9:0.1, *v/v*) or $MeOH/HCO_2H$ (99.9:0.1, *v/v*) as eluent mixture; the ionization source was electrospray in positive mode ($ESI^{+}$) with a voltage of 15 V; the range of masses in acquisition was 50–1200 u in SCAN mode. Flash-column chromatography was carried out in an automated system, using silica gel (230–400 mesh). Analytical thin layer chromatography (TLC) was performed on aluminum plates with Merck Kieselgel $60F_{254}$ and visualized by UV irradiation (254 nm) or by staining with a solution of phosphomolybdic acid, $KMnO_4$, or *p*-anisaldehyde. Melting points were measured in a Büchi B-540 apparatus in open capillary tubes.

**Ethyl (2-Iodophenyl)carbamate (2a)** (Denis et al., 2015)**.**

**Ethyl (2-Iodo-4-methylphenyl)carbamate (2b).**

**General procedure for aniline protection as carbamate**. To a cooled (0 ºC) suspension of $K_2CO_3$ (0.36 g, 2.58 mmol) and 2-iodo-4-methylaniline **1b** (0.50 g, 2.15 mmol) in THF (11 mL), was added dropwise ethyl chloroformate (0.23 mL, 2.36 mmol) and the mixture was stirred at room temperature for 3 h. The reaction was poured over $H_2O$ and the mixture was extracted with $CH_2Cl_2$ (3x). The combined organic layers were washed with a saturated aqueous solution of $NaHCO_3$ and dried with anhydrous $Na_2SO_4$. The solvent was evaporated, and the residue was adsorbed on silica gel before purification by flash-column chromatography (silica gel, gradient from 100:0 to 80:20 *v/v*, *n*-hexane/EtOAc) to afford 0.49 g (74% yield) of **2b**. **m.p.**: 52-54 °C (hexane/EtOAc). **$^{1}$H NMR** (400.13 MHz, $CDCl_3$): δ 7.90 (d, *J* = 8.4 Hz, 1H), 7.61 (dd, *J* = 2.0, 0.9 Hz, 1H), 7.16 (d, *J* = 8.4, 2.0 Hz, 1H), 6.84 (br s, 1H), 4.26 (q, *J* = 7.1 Hz, 2H), 2.29 (d, *J* = 0.7 Hz, 3H), 1.35 (t, *J* = 7.1 Hz, 3H) ppm. **$^{13}$C NMR** (100.61 MHz, $CDCl_3$): δ 153.6 (s), 139.0 (d), 136.0 (s), 135.0 (s), 129.3 (d), 120.3 (s), 89.1 (d), 61.5

(t), 20.2 (q), 14.5 (q) ppm. **IR** (NaCl): ν 3390 (w, N-H), 1736 (s, C=O), 1520 (s), 1220 (s, C-O) $cm^{-1}$. **HRMS** ($ESI^+$): Calcd for $C_{10}H_{13}INO_2$ ($[M+H]^+$), 305.9985; found, 305.9975.

**Ethyl (2-Iodo-4-methoxyphenyl)carbamate (2c).** Following the general procedure for aniline protection as carbamate described above, the reaction of $K_2CO_3$ (0.14 g, 1.04 mmol), 2-iodo-4-methoxyaniline **1c** (0.20 g, 0.80 mmol) and ethyl chloroformate (0.09 mL, 0.96 mmol) in THF (4 mL) afforded, after purification by flash-column chromatography (silica gel, gradient from 100:0 to 80:20 *v/v*, *n*-hexane/EtOAc), 0.21 g (81% yield) of **2c**. **m.p.**: 79-80 ºC (hexane/EtOAc). **$^1$H NMR** (400.13 MHz, $CDCl_3$): δ 7.81 (br d, *J* = 9.0 Hz, 1H), 7.30 (d, *J* = 2.9 Hz, 1H), 6.90 (dd, *J* = 9.0, 2.9 Hz, 1H), 6.65 (br s, 1H), 4.23 (q, *J* = 7.1 Hz, 2H), 3.76 (s, 3H), 1.32 (t, *J* = 7.1 Hz, 3H) ppm. **$^{13}$C NMR** (100.61 MHz, $CDCl_3$): δ 156.4 (s), 154.0 (s), 132.1 (s), 123.9 (d), 122.2 (d), 115.1 (d), 90.9 (s), 61.6 (t), 55.8 (q), 14.7 (q) ppm. **IR** (NaCl): ν 3279 (s, N-H), 2978 (w), 2938 (w), 2904 (w), 2838 (w), 1697 (s, C=O), 1528 (s), 1401 (w), 1283 (s, C-O-C), 1240 (s, C-O), 1216 (s, C-O), 1034 (w), 1024 (w), 854 (w) $cm^{-1}$. **HRMS** ($ESI^+$): Calcd for $C_{10}H_{13}INO_3$ ($[M+H]^+$), 321.9935; found, 321.9931.

**Ethyl (4-Bromo-2-iodophenyl)carbamate (2d).** Following the general procedure for aniline protection as carbamate described above, the reaction of 4-bromo-2-iodoaniline **1d** (0.50 g, 1.68 mmol), ethyl chloroformate (0.19 mL, 2.01 mmol) and $K_2CO_3$ (301.5 mg, 2.18 mmol) in THF (8.4 mL) afforded, after purification by flash-column chromatography (silica gel, gradient from 100:0 to 80:20 *v/v*, *n*-hexane/EtOAc), 0.55 g (88% yield) of **2d**. **m.p.**: 105-106 °C (*n*-hexane/EtOAc). **$^1$H NMR** (400.13 MHz, $CDCl_3$): δ 8.00 (d, *J* = 8.9 Hz, 1H), 7.90 (d, *J* = 2.3 Hz, 1H), 7.47 (dd, *J* = 8.9, 2.3 Hz, 1H), 6.93 (br s, 1H), 4.27 (q, *J* = 7.1 Hz, 2H), 1.36 (t, *J* = 7.1 Hz, 3H) ppm. **$^{13}$C NMR** (100.61 MHz, $CDCl_3$): δ 153.3 (s), 140.5 (d), 137.8 (s), 132.2 (d), 120.9 (d), 116.2 (s), 88.8 (s), 61.8 (t), 14.5 (q) ppm. **IR** (NaCl): ν 3294 (s, N-H), 2980 (w, C-H), 2956 (w, C-H), 2929 (w, C-H), 1690 (s, C=O), 1562 (s), 1518 (s), 1075 (s, C-Br) $cm^{-1}$. **HRMS** ($ESI^+$): Calcd for $C_9H_9{}^{79}BrINO_2$ ($[M+H]^+$), 368.8861; found, 368.8865; Calcd for $C_9H_9{}^{81}BrINO_2$ ($[M+H]^+$), 370.8841; found, 370.8845.

**Ethyl (4-Methoxycarbonyl-2-iodophenyl)carbamate (2e)** (Vaz et al., 2021)**.**

**Ethyl (*E*)-4-(2-Ethynylphenylamino)-4-oxobut-2-enoate (4a).**

**General procedure for aniline protection as amide**. To a suspension of 2-ethynylaniline **3a** (0.49 mL, 4.27 mmol) and $NaHCO_3$ (0.72 g, 8.54 mmol) in dioxane (8 mL) at 0 °C was added dropwise ethyl fumaroyl chloride (0.71 mL, 6.40 mmol). After 23 h at 25 ºC, the reaction was poured into water and the mixture was extracted with EtOAc (3x). The combined organic layers were washed with water and dried. The solvent was evaporated, and the residue was adsorbed on silica gel before purification by flash-column chromatography (silica gel, gradient from 100:0 to 80:20 *v/v*, *n*-hexane/EtOAc), to afford 0.94 g (90% yield) of **4a**. **m.p.**: 139 °C ($Et_2O$). **$^1$H NMR** (400.13 MHz, $CDCl_3$): δ 8.51 (d, *J* = 8.4 Hz, 1H), 8.19 (br s, 1H), 7.49 (dd, *J* = 7.8, 1.5 Hz, 1H), 7.40 (td, *J* = 8.0, 1.6 Hz, 1H), 7.10 (td, *J* = 7.6, 1.1 Hz, 1H), 7.07 (d, *J* = 15.3 Hz, 1H), 6.96 (d, *J* = 15.3 Hz, 1H), 4.29 (q, *J* = 7.1 Hz, 2H), 3.58 (s, 1H), 1.35 (t, *J* = 7.1 Hz, 3H) ppm. **$^{13}$C NMR** (100.61 MHz, $CDCl_3$): δ 165.4 (s), 161.5 (s), 139.0 (d), 136.4 (s), 132.4 (d), 131.9 (d), 130.4 (d), 124.3 (d), 119.7 (d), 111.2 (s), 85.3 (s), 79.0 (d), 61.5 (t), 14.2 (q) ppm. **IR** (NaCl): ν 3289 (w, N–H), 3247 (s, ≡C–H), 1712 (s, C=O),

1641 (s, C=O) $cm^{-1}$. **HRMS** ($ESI^+$): Calcd for $C_{14}H_{13}NO_3$ ($[M]^+$), 243.0895; found, 243.0893.

**Ethyl (*E*)-4-(2-Ethynyl-4-methylphenylamino)-4-oxobut-2-enoate (4b).** Following the general procedure for aniline protection as amide described above, the reaction of ethyl fumaroyl chloride (0.71 mL, 6.16 mmol), 2-ethynyl-4-methylaniline **3b** (0.34 mL, 4.11 mmol) and $NaHCO_3$ (690.4 mg, 8.22 mmol) in dioxane (14 mL) afforded, after purification by flash-column chromatography (silica gel, gradient from 100:0 to 80:20 *v/v*, *n*-hexane/EtOAc), 0.67 g (66% yield) of **4b**. **m.p.**: 138-140 °C (*n*-hexane/EtOAc). **$^1$H NMR** (400.13 MHz, $CDCl_3$): δ 8.38 (d, *J* = 8.5 Hz, 1H), 8.12 (br s, 1H), 7.30 (s, 1H), 7.20 (d, *J* = 8.0 Hz, 1H), 7.06 (d, *J* = 15.3 Hz, 1H), 6.94 (d, *J* = 15.3 Hz, 1H), 4.29 (q, *J* = 7.1 Hz, 2H), 3.53 (s, 1H), 2.30 (s, 3H), 1.34 (t, *J* = 7.1 Hz, 3H) ppm. **$^{13}$C NMR** (100.61 MHz, $CDCl_3$): δ 165.5 (s), 161.4 (s), 136.7 (s), 136.6 (d), 134.1 (s), 132.7 (d), 131.7 (d), 131.2 (d), 119.6 (d), 111.1 (s), 84.8 (s), 79.2 (d), 61.5 (t), 20.8 (q), 14.3 (q) ppm. **IR** (NaCl): ν 3280 (w, N–H), 3252 (s, C≡C-H), 2980 (w, C-H), 1711 (s, C=O), 1667 (w, C=O), 1642 (s, N-H), 1534 (w, N-H), 1298 (w, C-O-C), 675 (w) $cm^{-1}$. **HRMS** ($ESI^+$): Calcd. for $C_{15}H_{16}NO_3$ ($[M+H]^+$); 258.1125; found, 258.1132.

**Ethyl (*E*)-4-(2-Ethynyl-4-fluorophenylamino)-4-oxobut-2-enoate (4c).** Following the general procedure for aniline protection as amide described above, the reaction of ethyl fumaroyl chloride (0.39 mL, 3.40 mmol), 2-ethynyl-4-fluoroaniline **3c** (0.32 g, 2.28 mmol) and $NaHCO_3$ (0.38 g, 4.60 mmol) in dioxane (8 mL) afforded, after purification by flash-column chromatography (silica gel, gradient from 100:0 to 80:20 *v/v*, *n*-hexane/EtOAc), 0.43 g (72% yield) of a white solid identified as **4c**. **m.p.:** 170-172 °C (*n*-hexane/EtOAc). **$^1$H NMR** (400.13 MHz, $CDCl_3$): δ 8.50 (dd, *J* = 9.2 Hz, and $^4J_{H,F}$ = 5.2 Hz, 1H), 8.11 (br s, 1H), 7.21 (dd, *J* = 3.0 Hz, and $^3J_{H,F}$ = 8.4 Hz, 1H), 7.13 (ddd, *J* = 9.2, 3.0 Hz, and $^3J_{H,F}$ = 8.0 Hz, 1H), 7.08 (d, *J* = 15.3 Hz, 1H), 6.97 (d, *J* = 15.3 Hz, 1H), 4.31 (q, *J* = 7.1 Hz, 2H), 3.63 (s, 1H), 1.37 (t, *J* = 7.1 Hz, 3H) ppm. **$^{13}$C NMR** (100.61 MHz, $CDCl_3$): δ 165.3 (s), 161.3 (s), 158.5 (s, $^1J_{C,F}$ = 245.7 Hz), 136.1 (d), 135.4 (s, $^4J_{C,F}$ = 2.9 Hz), 132.0 (d), 121.5 (d, $^3J_{C,F}$ = 8.3 Hz), 118.7 (d, $^2J_{C,F}$ = 18.7 Hz), 117.5 (d, $^2J_{C,F}$ = 22.1 Hz), 112.8 (s, $^3J_{C,F}$ = 9.2 Hz), 85.9 (s), 77.9 (d, $^5J_{C,F}$ = 2.9 Hz), 61.5 (t), 14.1 (q) ppm. **IR** (NaCl): ν 3282 (w, N-H), 3245 (s, N-H), 3077 (w), 1711 (s, C=O), 1668 (w, C=O), 1644 (s), 1538 (w), 1302 (w, C-O-C), 674 (w) $cm^{-1}$. **HRMS** ($ESI^+$): Calcd. for $C_{14}H_{13}FNO_3$ ($[M+H]^+$), 262.0874; found, 262.0871.

**Ethyl (*Z*)-7-(2-Ethoxy-2-oxoethylidene)-dihydrobenzo[2,3]azepino[4,5-*b*]indole-12(5*H*)-carboxylate (5a, UVI3508)**

**General procedure for one-pot Sonogashira-Heterocyclization-oxidative Heck reaction**. To a degassed solution of ethyl (2-iodophenyl)carbamate **2a** (0.20 g, 0.69 mmol) and alkyne **4a** (0.33 g, 1.37 mmol) in DMF (7.6 mL) were sequentially added $Et_3N$ (0.43 mL, 3.09 mmol), $PdCl_2(PPh_3)$ (0.048 g, 0.069 mmol), CuI (0.052 g, 0.28 mmol) and $PPh_3$ (0.018 g, 0.069 mmol). The mixture was stirred at 60 °C for 12 h. Then, the flask was opened to air and the reaction mixture was stirred at 100 °C for 15 h. A water/brine solution (90:10 *v/v*) was added and the mixture was extracted with EtOAc (3x). The combined organic layers were dried. The solvent was evaporated, and the residue was adsorbed on silica gel before purification by column chromatography (silica gel, gradient from 100:0 to 60:40 *v/v*, hexane/EtOAc) to afford 0.15 g (52% yield) of **5a**.[6] **$^1$H NMR** (400.13 MHz, DMSO-$d_6$, 333 K): δ 8.83 (s, 1H), 8.20 (d, *J* = 8.3 Hz, 1H), 7.79 (dd, *J* = 7.7, 1.2 Hz, 1H), 7.45 (ddd, *J* = 8.5, 7.3, 1.3 Hz, 1H), 7.41 – 7.31 (m,

3H), 7.28 (d, *J* = 7.9 Hz, 1H), 7.20 (td, *J* = 7.5, 1.4 Hz, 1H), 6.31 (s, 1H), 4.35 (q, *J* = 7.2 Hz, 2H), 4.25 (q, *J* = 7.1 Hz, 2H), 1.31 (t, *J* = 7.1 Hz, 3H), 1.21 (t, *J* = 7.1 Hz, 3H) ppm. $^{13}$**C NMR** (100.61 MHz, DMSO-$d_6$, 333 K): δ 168.9 (s), 164.9 (s), 151.7 (s), 139.2 (s), 138.8 (s), 134.0 (s), 133.1 (s), 130.1 (d), 128.9 (d), 126.6 (d), 126.6 (s), 125.2 (d), 124.3 (d), 124.1 (d), 124.1 (s), 123.0 (d), 121.9 (s), 119.3 (d), 115.6 (d), 63.9 (t), 61.3 (t), 14.1 (q), 14.0 (q) ppm.

**Ethyl (*Z*)-7-(2-Ethoxy-2-oxoethylidene)-3-methyl-6-oxo-6,7-dihydrobenzo[2,3]azepino[4,5-*b*]indole-12(5*H*)-carboxylate (5b, UVI3511).** Following the general procedure for the one-pot Sonogashira-heterocyclization-oxidative Heck reaction described above, the reaction of iodide **2a** (0.20 g, 0.69 mmol), alkyne **4b** (0.35 g, 1.37 mmol), $Et_3N$ (0.43 mL, 3.09 mmol), $PdCl_2(PPh_3)_2$ (0.048 g, 0.069 mmol), CuI (0.052 g, 0.28 mmol) and $PPh_3$ (0.018 g, 0.069 mmol) in DMF (7.6 mL) afforded, after purification by flash-column chromatography (silica gel, gradient from 100:0 to 60:40 *v/v*, *n*-hexane/EtOAc), 0.22 g (75% yield) of **5b**. **m.p.:** 255-257 °C (*n*-hexane/EtOAc). $^{1}$**H NMR** (400.13 MHz, $CDCl_3$, 323 K): δ 8.67 (s, 1H), 8.20 (d, *J* = 8.3 Hz, 1H), 7.78 (dt, *J* = 7.8, 1.1 Hz, 1H), 7.43 (ddd, *J* = 8.4, 7.2, 1.3 Hz, 1H), 7.35 (t, *J* = 7.5 Hz, 1H), 7.19 – 7.12 (m, 2H), 7.15 (s, 1H), 6.28 (s, 1H), 4.35 (q, *J* = 7.1 Hz, 2H), 4.25 (q, *J* = 7.1 Hz, 2H), 2.36 (s, 3H), 1.30 (t, *J* = 7.1 Hz, 3H), 1.21 (t, *J* = 7.1 Hz, 3H) ppm. $^{13}$**C NMR** (100.61 MHz, $CDCl_3$, 323 K): δ 168.9 (s), 164.9 (s), 151.7 (s), 139.4 (s), 137.1 (s), 134.2 (s), 133.6 (s), 130.9 (s), 130.2 (d), 129.8 (d), 126.7 (s), 126.5 (d), 125.2 (d), 124.2 (d), 124.1 (s), 122.9 (d), 121.9 (s), 119.3 (d), 115.5 (d), 63.8 (t), 61.2 (t), 21.4 (q), 14.1 (q), 14.0 (q) ppm. **IR** (NaCl): ν 3196 (w, N-H), 2981(s, C-H), 2930 (w, C-H), 1731 (s, C=O), 1666 (s, C=O), 1585 (w, N-C=O), 1505 (s), 1454 (w, $-CH_3$), 1371 (s), 1307 (s), 1226 (s, C-O-C), 759 (s) $cm^{-1}$. **HRMS** ($ESI^+$): Calcd. for $C_{24}H_{23}N_2O_5$ ($[M+H]^+$), 419.1601; found, 419.1588.

**Ethyl (*Z*)-7-(2-Ethoxy-2-oxoethylidene)-3-methyl-9-bromo-6-oxo-6,7-dihydrobenzo[2,3]azepino[4,5-*b*]indole-12(5*H*)-carboxylate (5c, UVI3507).** Following the general procedure for the one-pot Sonogashira-heterocyclization-oxidative Heck reaction described above, the reaction of iodide **2d** (0.15 g, 0.41 mmol), alkyne **4b** (0.21 g, 0.81 mmol), $Et_3N$ (0.25 mL, 1.82 mmol), $PdCl_2(PPh_3)_2$ (29 mg, 0.041 mmol, CuI (31 mg, 0.031 mmol) and $PPh_3$ (11 mg, 0.041 mmol) in DMF (4.5 mL) afforded, after purification by flash-column chromatography (silica gel, gradient from 100:0 to 60:40 *v/v*, *n*-hexane/EtOAc), 0.11 g (55% yield) of **5c**. **m.p.:** 271-274 °C (*n*-hexane/EtOAc). $^{1}$**H NMR** (400.13 MHz, $CDCl_3$, 298 K): δ 8.66 (br s, 1H), 8.10 (d, *J* = 8.9 Hz, 1H), 7.88 (d, *J* = 2.0 Hz, 1H), 7.52 (dd, *J* = 8.9, 2.0 Hz, 1H), 7.18 – 7.16 (m, 2H), 7.13 (s, 1H), 6.27 (s, 1H), 4.40 – 4.29 (m, 2H), 4.25 (q, *J* = 7.1 Hz, 2H), 2.35 (s, 3H), 1.31 (t, *J* = 7.1 Hz, 3H), 1.20 (t, *J* = 7.1 Hz, 3H) ppm. $^{13}$**C NMR** (100.61 MHz, $CDCl_3$, 298 K): δ 168.6 (s), 164.7 (s), 151.3 (s), 138.9 (s), 137.4 (s), 135.2 (s), 133.8 (s), 130.9 (s), 130.3 (d), 130.2 (d), 129.3 (d), 128.1 (s), 125.4 (d), 123.5 (s), 123.1 (d), 121.9 (d), 120.6 (s), 117.6 (s), 117.0 (d), 64.0 (t), 61.2 (t), 20.9 (q), 14.0 (q), 13.8 (q) ppm. **IR** (NaCl): ν 2980 (s, C-H), 2932 (w, C-H), 1731 (s, C=O), 1666 (s, C=O), 1595 (w, N-C=O), 1505 (s), 1449 (w), 1372 (s), 1262 (s), 1227 (s), 1176 (s), 1043 (s) $cm^{-1}$. **HRMS** ($ESI^+$): Calcd. for $C_{24}H_{22}{}^{79}BrN_2O_5$ ($[M+H]^+$), 497.0707; found, 497.0691.

**Ethyl (*Z*)-7-(2-Ethoxy-2-oxoethylidene)-2-fluoro-9-methyl-6-oxo-6,7-dihydrobenzo[2,3]azepino[4.5-b]indole-12(5*H*)-carboxylate (5d, UVI3503).** Following the general procedure for the one-pot Sonogashira-heterocyclization-oxidative Heck reaction described above, the reaction of iodide **2b** (0.15 g, 0.49 mmol), alkyne **4c** (0.26 g, 0.98 mmol), $Et_3N$ (0.31 mL, 2.21 mmol), $PdCl_2(PPh_3)_2$ (34 mg, 0.049 mmol), CuI (38 mg, 0.20 mmol) and $PPh_3$ (13 mg, 0.049 mmol) in DMF (5.5 mL) afforded, after purification by flash-column chromatography (silica gel, gradient from 100:0 to 60:40 *v/v*, *n*-hexane/EtOAc), 0.1 g (47% yield) of **5d**. **m.p.:** 226-227 °C (MeOH). **$^1$H NMR** (400.13 MHz, $CDCl_3$, 323 K): δ 9.27 (s, 1H), 8.08 (d, *J* = 8.5 Hz, 1H), 7.56 (s, 1H), 7.33 – 7.27 (m, 2H), 7.08 – 6.99 (m, 2H), 6.29 (s, 1H), 4.38 (q, *J* = 7.1 Hz, 2H), 4.24 (q, *J* = 7.1 Hz, 2H), 2.48 (s, 3H), 1.30 (t, *J* = 7.1 Hz, 3H), 1.26 (t, *J* = 7.1 Hz, 3H) ppm. **$^{13}$C NMR** (100.61 MHz, $CDCl_3$, 323 K): δ 169.2 (s), 164.8 (s), 158.9 (s, $^1J_{C,F}$ = 244.1 Hz), 151.4 (s), 139.3 (s), 137.2 (s), 134.2 (s), 133.0 (s, $^4J_{C,F}$ = 1.5 Hz), 129.6 (s, $^4J_{C,F}$ = 2.7 Hz), 128.4 (d), 126.7 (s), 126.0 (s, $^3J_{C,F}$ = 9.1 Hz), 126.3 (d), 124.9 (d, $^3J_{C,F}$ = 8.5 Hz), 122.7 (s), 119.2 (d), 116.2 (d, $^2J_{C,F}$ = 20.6 Hz), 115.9 (d, $^2J_{C,F}$ = 19.0 Hz), 115.5 (d), 63.9 (t), 61.3 (t), 21.4 (q), 14.1 (q), 14.0 (q) ppm. **IR** (NaCl): ν 3199 (w, N-H), 3074 (w, N-H), 2980 (w, C-H), 2938 (w, C-H), 1727 (s, C=O), 1666 (s, C=O), 1502 (m), 1396 (m), 1307 (m), 1185 (s), 1041 (m) $cm^{-1}$. **HRMS** ($ESI^+$): Calcd. for $C_{24}H_{22}FN_2O_5$ ($[M+H]^+$); 437.1507; found, 437.1508.

**12-Ethyl 9-Methyl (*Z*)-7-(2-Ethoxy-2-oxoethylidene)-2-methyl-6-oxo-6,7-dihydrobenzo[2,3]azepino[4,5-b]indole-12, 9(5*H*)-dicarboxylate (5e, UVI3501).** Following the general procedure for the one-pot Sonogashira-heterocyclization-oxidative Heck reaction described above, the reaction of iodide **2e** (0.15 g, 0.43 mmol), alkyne **x.x** (0.22 g, 0.86 mmol), $Et_3N$ (0.27 mL, 1.94 mmol), $PdCl_2(PPh_3)_2$ (30 mg, 0.043 mmol), CuI (33 mg, 0.17 mmol) and $PPh_3$ (11 mg, 0.043 mmol) in DMF (4.8 mL) afforded, after purification by flash-column chromatography (silica gel, gradient from 100:0 to 60:40 *v/v*, *n*-hexane/EtOAc), 0.11 g (51% yield) of **5e**. **m.p.:** 263-264 °C ($CHCl_3$). **$^1$H NMR** (400.13 MHz, $CDCl_3$, 323 K): δ 8.66 (s, 1H), 8.47 (d, *J* = 1.7 Hz, 1H), 8.23 (d, *J* = 8.8 Hz, 1H), 8.13 (dd, *J* = 8.8, 1.7 Hz, 1H), 7.19 – 7.17 (m, 2H), 6.30 (s, 1H), 4.36 (q, *J* = 7.1 Hz, 2H), 4.26 (q, *J* = 7.1 Hz, 2H), 3.97 (s, 3H), 2.36 (s, 3H), 1.32 (t, *J* = 7.1 Hz, 3H), 1.21 (t, *J* = 7.1 Hz, 3H) ppm. **$^{13}$C NMR** (100.61 MHz, $CDCl_3$, 323 K): δ 168.8 (s), 167.0 (s), 164.7 (s), 151.3 (s), 141.4 (s), 139.0 (s), 135.4 (s), 133.8 (s), 131.2 (s), 130.2 (d), 130.1 (d), 127.7 (d), 126.6 (s), 126.5 (s), 125.7 (d), 123.6 (s), 123.1 (d), 121.7 (s), 121.4 (d), 115.2 (d), 64.2 (t), 61.3 (t), 52.3 (q), 21.0 (q), 14.1 (q), 13.9 (q) ppm. **IR** (NaCl): ν 3278 (w, N-H), 3031 (w, C-H), 2995 (w, C-H), 1721 (s, C=O), 1665 (s, C=O), 1373 (m), 1263 (s), 1226 (s), 1079 (m) $cm^{-1}$. **HRMS** ($ESI^+$): Calcd. for $C_{26}H_{25}N_2O_7$ ($[M+H]^+$); 477.1656; found, 477.1656.

**Ethyl (*Z*)-7-(2-Ethoxy-2-oxoethylidene)-9-methoxy-2-methyl-6-oxo-6,7-dihydrobenzo[2,3]azepino[4,5-*b*]indole-12(5*H*)-carboxylate (5f, UVI3502).** Following the general procedure for the one-pot Sonogashira-heterocyclization-oxidative Heck reaction described above, the reaction of iodide **x.x** (0.15 g, 0.47 mmol), alkyne **x.x** (0.24 g, 0.93 mmol), $Et_3N$ (0.29 mL, 2.10 mmol), $PdCl_2(PPh_3)_2$ (33 mg, 0.047 mmol), CuI (36 mg, 0.19 mmol) and $PPh_3$ (12 mg, 0.047 mmol) in DMF (5.2 mL) afforded, after purification by flash-column chromatography (silica gel, gradient from 100:0 to 60:40 *v/v*, *n*-hexane/EtOAc), 0.12 g (55% yield) of **5f**. **m.p.:** 159-160 °C (MeOH). **$^1$H NMR** (400.13 MHz, $CDCl_3$, 323 K): δ 9.35 (s, 1H), 8.08 (d, *J* = 9.1 Hz, 1H), 7.24 – 7.18 (m, 2H), 7.16 – 7.08 (m, 2H), 7.03 (dd, *J* = 9.1, 2.5 Hz, 1H), 6.25 (s, 1H), 4.32 (q, *J* = 7.1 Hz, 2H), 4.23 (q, *J* = 7.1 Hz, 2H), 3.88 (s, 3H), 2.34 (s, 3H), 1.28 (t, *J* = 7.1 Hz, 3H), 1.18 (t, *J* = 7.1 Hz, 3H) ppm. **$^{13}$C NMR** (100.61 MHz, $CDCl_3$, 323 K): δ 169.2 (s), 164.8 (s), 157.2 (s), 151.6 (s), 139.8 (s), 134.8 (s), 133.5 (s), 133.4 (s), 131.0

(s), 130.2 (d), 129.7 (d), 127.4 (s), 124.6 (d), 123.9 (s), 123.0 (d), 121.6 (s), 116.4 (d), 115.4 (d), 101.7 (d), 63.6 (t), 61.1 (t), 55.9 (q), 20.9 (q), 14.0 (q), 13.9 (q) ppm. **IR** (NaCl): ν 3289 (w, N-H), 3194 (w, N-H), 2980 (w, C-H), 2938 (w, C-H), 2835 (w, C-H), 1731 (s, C=O), 1664 (s, C=O), 1506 (m), 1480 (m), 1373 (m), 1252 (m), 1163 (m), 1031 (m), 772 (s) $cm^{-1}$. **HRMS** ($ESI^+$): Calcd. for $C_{25}H_{25}N_2O_6$ ($[M+H]^+$); 449.1707; found, 449.1701.

**Ethyl 2-(6-Oxo-5,6,7,12-tetrahydrobenzo[2,3]azepino[4,5-*b*]indol-7-yl)acetate (7a, UVI3505).** To a solution of benzoazepinone **5a** (0.23 g, 0.57 mmol) in EtOAc (38 mL) was added $Pd(OH)_2$ (0.42 g, 0.60 mmol) and the resulting mixture was stirred at 25 °C under $H_2$ atmosphere (previous cycles of vacuum/$H_2$ were performed) for 24 h. After completion, the reaction mixture was filtered through a pad of Celite® and the product was eluted with 80:20 *v/v* $CH_2Cl_2$/MeOH. The product thus obtained was used in the next step without further purification. The solid obtained was introduced in a Schlenk tube and dissolved in THF (6.4 mL). A solution of TBAF (1M in THF, 2.87 mL, 2.87 mmol) was added and the mixture was stirred at 80 °C for 21 h. Then, an aqueous saturated solution of $NH_4Cl$ was added and the mixture was extracted with EtOAc (3x). The combined organic layers were dried over $Na_2SO_4$, the solvent was evaporated, and the residue was adsorbed on silica gel before purification by flash-column chromatography (silica gel, gradient from 100:0 to 90:10 *v/v*, $CH_2Cl_2$/MeOH) to afford 0.14 g (73% yield, two steps) of a yellow solid identified as **7a**. **m.p.:** 257-259 °C ($CH_2Cl_2$/MeOH). **$^1$H NMR** (400.13 MHz, DMSO-$d_6$, 333 K): δ 10.08 (s, 1H), 7.79 (d, *J* = 8.0 Hz, 1H), 7.66 (d, *J* = 8.0 Hz, 1H), 7.44 (d, *J* = 8.0 Hz, 1H), 7.43 – 7.37 (m, 1H), 7.33 – 7.27 (m, 2H), 7.17 (dd, *J* = 8.2, 7.0 Hz, 1H), 7.07 (t, *J* = 7.5 Hz, 1H), 4.05 – 3.92 (m, 3H), 3.06 (dd, *J* = 15.6, 8.5 Hz, 1H), 2.73 (dd, *J* = 15.6, 7.2 Hz, 1H), 1.10 (t, *J* = 7.1 Hz, 3H) ppm. **$^{13}$C NMR** (100.63 MHz, DMSO-$d_6$, 333 K): δ 170.8 (s), 170.4 (s), 137.3 (s), 134.9 (s), 132.1 (s), 127.9 (d), 126.6 (d), 125.8 (s), 123.3 (d), 122.3 (s), 121.7 (d), 121.5 (d), 118.9 (d), 118.2 (d), 111.3 (d), 109.7 (s), 59.6 (t), 39.5 (d), 33.0 (t), 13.6 (q) ppm. **IR** (NaCl): ν 3301 (w, N-H), 3048 (w), 2970 (w), 1722 (s, C=O), 1645 (s, NHC=O), 1183 (s, C-O-C) $cm^{-1}$. **HRMS** ($ESI^+$): Calcd. for $C_{20}H_{19}N_2O_3$ ($[M+H]^+$); 335.1390; found, 335.1392.

**Ethyl 2-(6-Methyl-9,14-dihydrobenzo[2,3][1,2,4]triazolo[4',3':1,7]azepino[4,5-*b*]indol-9-yl)acetate (9, UVI8113).** Into a round-bottomed flask was added benzoazepinone **7a** (0.13 g, 0.39 mmol), Lawesson's reagent (0.32 g, 0.79 mmol) and THF (5.8 mL). The mixture was stirred at 60 °C for 3 h. The mixture was then diluted with $CH_2Cl_2$ and a saturated aqueous solution of $NaHCO_3$ was added. After extraction with $CH_2Cl_2$ (3x), the combined organic layers were washed with water (1x) and brine (1x). The solution was then dried over anhydrous $Na_2SO_4$, filtered, and concentrated. The residue was adsorbed on silica gel before purification by flash-column chromatography (silica gel, gradient from 100:0 to 95:5 *v/v*, $CH_2Cl_2$/MeOH) to afford 0.13 g (94%) of the thioacetamide, which was directly used in the next transformation.

Hydrazine hydrate was added dropwise to a stirred solution of the thioacetamide (22 mg, 0.062 mmol) in THF (0.3 mL) at 20 °C. After 2 h, the solution was concentrated under reduced pressure and the residue was treated with triethyl orthoacetate (0.40 mL, 2.17 mmol). The mixture was heated at 80 °C for 1 h. After cooling down to room temperature, a saturated aqueous solution of $NH_4Cl$ and EtOAc were added, and the mixture was extracted with EtOAc (3x). The combined organic layers were dried over $Na_2SO_4$ and concentrated under reduced pressure. The residue was dissolved in the minimum amount of $CH_2Cl_2$ and $Et_2O$ was added to form a precipitate. The solid was washed with $Et_2O$ and dried to afford 20.2 mg (87% yield) of **9**. **$^{1}$H NMR** (400.13 MHz, Methanol-*d*4, 333 K): δ 7.91 (dd, *J* = 7.8, 1.6 Hz, 1H), 7.74 (d, *J* = 8.4 Hz, 1H), 7.71 (dd, *J* = 8.1, 1.1 Hz, 2H), 7.65 (td, *J* = 7.6, 1.5 Hz, 1H), 7.59 (td, *J* = 7.7, 1.6 Hz, 1H), 7.43 (dt, *J* = 8.1, 0.9 Hz, 1H), 7.20 (t, *J* = 7.5 Hz, 1H), 7.12 (t, *J* = 7.6 Hz, 1H), 5.33 (t, *J* = 8.2 Hz, 1H), 3.90 (q, *J* = 6.5 Hz, 2H), 2.56 (dd, *J* = 14.7, 7.1 Hz, 1H), 2.49 (s, 3H), 2.42 (dd, *J* = 14.7, 8.5 Hz, 1H), 1.02 (t, *J* = 7.3 Hz, 3H) ppm. **$^{13}$C NMR** (100.61 MHz, Methanol-*d*4, 333 K): δ 171.8 (s), 159.1 (s), 154.2 (s), 138.8 (s), 131.5 (s), 130.1 (d), 129.5 (d), 129.1 (d), 128.0 (s), 127.6 (s), 127.2 (s), 126.7 (d), 124.4 (d), 121.0 (d), 119.3 (d), 116.5 (s), 112.6 (d), 61.9 (t), 37.3 (t), 31.9 (d), 14.2 (q), 12.3 (q) ppm. **IR** (NaCl): ν 3180 (w), 3108 (w), 3065 (w), 3023 (w), 2981 (w), 1733 (s, C=O), 1532 (m), 1496 (m), 1457 (m), 1414 (m), 1326 (m), 1264 (m), 1181 (m), 1031 (m) $cm^{-1}$. **HRMS** ($ESI^+$): Calcd. for $C_{22}H_{21}N_4O_2$ ($[M+H]^+$); 373.1659; found, 373.1665.

**Ethyl (*Z*)-7-(2-Ethoxy-2-oxoethylidene)-2-methyl-6-oxo-9-phenyl-6,7-dihydrobenzo[2,3]azepino[4,5-b]-12(5*H*)-carboxylate (5h, UVI3510).** A reaction flask was charged with **5c** (20 mg, 0.04 mmol), phenylboronic acid (7.4 mg, 0.06 mmol), $Na_2CO_3$ (8.5 mg, 0.08 mmol), $Pd(OAc)_2$ (1 mg, 0.002 mmol) and P(*o*-tolyl)$_3$ (1.2 mg, 0.004 mmol) in 1,4-dioxane (0.25 mL) and $H_2O$ (0.04 mL). The mixture was degassed and stirred at 80 °C for 12 h. The crude mixture was filtered through a pad of silica gel and concentrated under reduced pressure. The crude product was purified by flash-column chromatography (silica gel, gradient from 80:20 to 50:50 *v/v*, *n*-hexane/EtOAc) to afford 10 mg (52% yield) of **5h**. **m.p.:** 136 – 138 °C ($CHCl_3$). **$^{1}$H NMR** (400.13 MHz, $CDCl_3$, 323K): δ 8.68 (s, 1H, NH), 8.26 (dd, *J* = 8.7 Hz, 1H), 7.95 (d, *J* = 1.2 Hz, 1H), 7.69 – 7.66 (m, 3H), 7.51 – 7.45 (m, 2H), 7.41 – 7.36 (m, 1H), 7.17 – 7.15 (m, 3H), 6.30 (s, 1H), 4.37 (brs, 2H), 4.24 (q, *J* = 7.1 Hz, 2H), 2.36 (s, 3H), 1.29 (t, *J* = 7.1 Hz, 3H), 1.22 (t, *J* = 7.1 Hz, 3H) ppm. **$^{13}$C NMR** (100.61 MHz, $CDCl_3$, 323K): δ 168.9 (s), 164.9 (s), 151.7 (s), 141.3 (s), 139.6 (s), 138.3 (s), 138.0 (s), 134.8 (s), 133.6 (s), 131.0 (s), 130.3 (d), 129.9 (d), 129.1 (d, 2x), 127.6 (d, 2x), 127.4 (d), 127.2 (s), 126.0 (d), 125.2 (d), 124.0 (s), 123.0 (d), 122.0 (s), 117.8 (d), 115.8 (d), 63.9 (t), 61.2 (t), 21.0 (q), 14.1 (q), 14.0 (q) ppm. **IR** (NaCl): ν 3189 (w), 3060 (w), 2980 (w), 2936 (w), 1727 (s, C=O), 1664 (s, C=O), 1465 (s), 1372 (s), 1228 (s) $cm^{-1}$. **HRMS** ($ESI^+$): Calcd. for $C_{30}H_{27}N_2O_5$ ($[M+H]^+$); 495.1914; found, 495.1917.

**(*Z*)-2-(6-Oxo-5,12-dihydrobenzo[2,3]azepino[4,5-b]indol-7(6*H*)-ylidene)acetic Acid (6a, UVI3506).** To a solution of **5a** (32 mg, 0.096 mmol) in THF (1.2 ml), a solution of LiOH $H_2O$ (80.3 mg, 1.91 mmol) in $H_2O$ (1.2 mL) was added and the mixture was stirred at 50 °C for 12 h. The reaction mixture was slowly quenched with $KHSO_4$ (1.0 M in water) to reach pH 4. Then, the mixture was extracted with EtOAc (3x) and the combined organic layers were washed with brine. The crude product was purified by flash-column chromatography (silica gel, gradient from 100:0 to 90:10 *v/v*, $CH_2Cl_2$/MeOH) to afford 25 mg (86% yield) of **6a**. **m.p.:** 258 – 260 °C (MeOH). **$^{1}$H NMR** (400.13 MHz, DMSO-*d*6): δ 12.62 (s, 1H), 12.33 (s, 1H), 10.66 (s, 1H), 7.87 (dd, *J* = 7.9, 1.5 Hz, 1H), 7.72 (d, *J* = 8.0 Hz, 1H), 7.54 (d, *J* = 8.1 Hz, 1H), 7.40 (ddd, *J* = 8.5, 7.1, 1.5 Hz, 1H), 7.34 – 7.22 (m, 3H), 7.16 (ddd, *J* = 8.0, 7.0, 1.1 Hz, 1H), 6.19 (s, 1H) ppm. **$^{13}$C NMR** (100.61 MHz, DMSO-*d*6): δ 166.9 (s), 166.4 (s), 138.6 (s), 137.9 (s), 134.3 (s), 133.5 (s), 128.7 (d), 127.3 (d), 125.2 (s), 123.9 (d), 123.2 (d), 122.9 (d), 122.1 (d), 121.5 (s), 120.4 (d), 118.3 (d), 112.0 (d), 110.6 (s) ppm. **IR** (NaCl): ν 3444 (s,

O-H, N-H), 2997(s), 3913 (s), 1661 (s, C=O), 1437 (m), 1407(m) $cm^{-1}$. **HRMS** ($ESI^+$): Calcd. for $C_{18}H_{13}N_2O_3$ ($[M+H]^+$); 305.0920; found, 305.0921.

**2-(6-Oxo-5,6,7,12-tetrahydrobenzo[2,3]azepino[4,5-b]indol-7-yl)acetic Acid (8, UVI3504).** Following the general reaction for ester hydrolysis described above, the reaction of **5a** (14 mg, 0.040 mmol), LiOH $H_2O$ (34 mg, 0.81 mmol) in THF (0.5 mL) and $H_2O$ (0.5 mL) afforded, after purification by flash-column chromatography (silica gel, gradient from 100:0 to 90:10 *v/v*, $CH_2Cl_2$/MeOH), 10 mg (79% yield) of **8**. **$^1$H NMR** (400.13 MHz, DMSO-$d_6$): δ 10.55 (s, 1H), 7.79 (dd, *J* = 7.8, 1.5 Hz, 1H), 7.65 – 7.59 (m, 2H), 7.49 -7.40 (m, 1H), 7.29 (td, *J* = 7.6, 1.3 Hz, 1H), 7.26 – 7.20 (m, 2H), 7.11 (td, *J* = 7.5, 0.9 Hz, 1H), 6.88 (s, 1H), 5.68 (dd, *J* = 10.1, 4.9 Hz, 1H), 2.86 (dd. *J* = 16.0, 4.9 Hz, 1H), 2.43 (dd, *J* = 10.1, 6.0 Hz, 1H) ppm. **$^{13}$C NMR** (100.61 MHz, DMSO-$d_6$): δ 170.2 (s), 167.8 (s), 136.6 (s), 135.5 (s), 134.0 (s), 129.5 (s), 129.4 (d), 129.1 (d), 127.7 (s), 124.9 (d), 123.1 (s), 121.9 (d), 121.4 (d), 120.3 (d), 109.9 (d), 101.7 (d), 57.4 (d), 34.7 (t) ppm. **IR** (NaCl): ν 3600-3000 (s, OH + NH), 3000.7 (m), 2916 (m), 1661 (s, C=O), 1437 (s), 1407 (s), 1314 (w) $cm^{-1}$. **HRMS** ($ESI^+$): Calcd. for $C_{18}H_{15}N_2O_3$ ($[M+H]^+$); 307.1077; found, 307.1078.

**(*Z*)-2-(9-Methoxy-2-methyl-6-oxo-5,12-dihydrobenzo[2,3]azepino[4,5-b]indol-7(6*H*)-ylidene)acetic Acid (6f, UVI3509).** To a solution of **5f** (40 mg, 0.089 mmol) in THF (1 mL), TBAF (0.45 mL, 0.45 mmol) was added. The resulting mixture was stirred at 80 °C for 21 h. Then, a saturated aqueous solution of $NH_4Cl$ was added and extracted with EtOAc (3x). The combined organic layers were dried over anhydrous $Na_2SO_4$ and concentrated. The residue was purified by flash-column chromatography (silica gel, gradient from 100:0 to 90:10 *v/v*, $CH_2Cl_2$/MeOH) to afford 22 mg (71% yield) of **6f**. **m.p.:** 293 – 295 °C (MeOH). **$^1$H NMR** (400.13 MHz, DMSO-$d_6$): δ 12.65 (s, 1H), 8.21 (s, 1H), 7.66 (d, *J* = 8.1 Hz, 1H), 7.55 (d, *J* = 8.7 Hz, 1H), 7.50 (dd, *J* = 8.3, 1.8 Hz, 1H), 7.45 (d, *J* = 2.3 Hz, 1H), 7.10 (dd, *J* = 8.7, 2.3 Hz, 1H) 6.97 (s, 1H), 3.91 (s, 3H), 2.52 (s, 3H) ppm. **$^{13}$C NMR** (100.61 MHz, DMSO-$d_6$): δ 168.6 (s), 155.7 (s), 154.8 (s), 140.4 (s), 139.1 (s), 139.0 (s), 136.6 (s), 134.4 (d), 133.1 (s), 131.8 (d), 126.2 (d), 125.7 (s), 123.4 (s), 116.3 (d), 113.2 (d), 108.5 (s), 103.9 (d), 103.2 (d), 55.9 (q), 21.1 (q) ppm. **IR** (NaCl): ν 3600-3000 (s, OH + N-H), 2999 (m), 2915 (m), 1662 (s, C=O), 1437 (w), 1407 (w) $cm^{-1}$. **HRMS** ($ESI^+$): Calcd. for $C_{20}H_{17}N_2O_4$ ($[M+H]^+$); 349.1183; found, 349.1184.

**Table S1.** $IC_{50}$ and $R^2$ values from the inhibition curves performed with the 12 compounds using $[^3H]$CP55,940 *vs.* increasing concentrations ranging from $10^{-12}$ M to $10^{-4}$ M of each compound in membrane homogenates from rat brain cortex.

| **Compound** | **$IC_{50}$ (nM)** | **$R^2$** |
|---|---|---|
| UVI3501 | 0.0005 ± >10000 | 0.2131 |
| UVI3502 | 0.47 ± 1.94<br>1470 ± 1.80 | 0.8071 |
| UVI3503 | >10000 | 0.1394 |
| UVI3504 | 5908 ± 7.18 | 0.3139 |
| UVI3505 | 8.85 ± 14.62 | 0.2335 |
| UVI3506 | 7041 ± 17.98 | 0.1755 |
| UVI3507 | 0.04 ± 21.03 | 0.2038 |
| UVI3508 | 3.1e5 ± 101.65 | 0.6952 |
| UVI3509 | >10000 | 0.1694 |
| UVI3510 | 1147 ± 1.65 | 0.8927 |
| UVI3511 | 0.03 ± 19.23 | 0.4512 |
| UVI8113 | 10.62 ± 43.35 | 0.1301 |

**Table S2.** [$^3$H]CP55,940 binding in different brain areas of rats (n = 5) and mice (n = 5) showing the total binding, [$^3$H]CP55,940 binding in the presence of UVI3502 and [$^3$H]CP55,940 binding in the presence of SR141716A.

| | [$^3$H]CP55,940 binding (fmol/mg t.e.) | | | | | |
|---|---|---|---|---|---|---|
| | **Sprague-Dawley rats** | | | **Swiss mice** | | |
| **Brain region** | **Total** | **UVI3502** | **SR141716A** | **Total** | **UVI3502** | **SR141716A** |
| Amygdala | 158 ± 28 | 136 ± 52 | 79 ± 18$^{##}$ | 168 ± 41 | 154 ± 8 | 62 ± 37$^{##}$ |
| Cerebellum (gray matter) | 724 ± 111 | 421 ± 126$^{*}$ | 72 ± 23$^{##}$ | 409 ± 115 | 409 ± 207 | 118 ± 22$^{##}$ |
| Cortex | 133 ± 28 | 122 ± 25 | 66 ± 14$^{#}$ | 81 ± 3 | 118 ± 18 | 65 ± 26 |
| Hippocampus dorsal | 299 ± 25 | 184 ± 26$^{*}$ | 62 ± 10$^{##}$ | 168 ± 15 | 181 ± 9 | 69 ± 29$^{#}$ |
| *Globus pallidus* | 1632 ± 65 | 572 ± 40$^{*}$ | 77 ± 15$^{#}$ | 1074 ± 262 | 698 ± 16$^{**}$ | 75 ± 31$^{##}$ |
| *Nucleus basalis magnocellularis* | 180 ± 19 | 147 ± 18 | 80 ± 11$^{##}$ | 237 ± 20 | 202 ± 19$^{*}$ | 82 ± 31$^{##}$ |
| Striatum | 560 ± 22 | 291 ± 18$^{*}$ | 79 ± 25$^{##}$ | 133 ± 15 | 159 ± 11 | 73 ± 13$^{##}$ |
| *Substantia nigra* | 2720 ± 540 | 1364 ± 581$^{*}$ | 70 ± 14$^{##}$ | 1684 ± 387 | 1075 ± 22$^{*}$ | 106 ± 118$^{##}$ |

Total *vs*. UVI3502 (*), Total *vs*. SR141716A ($^{#}$). Kruskal–Wallis test, *post-hoc* test Dunn's multiple comparison, $^{*#}p < 0.05$; $^{**##}p < 0.01$.

**Table S3.** [$^{35}$S]GTPγS binding in different brain areas of rats (n = 5) and mice (n = 5) in the presence of CP55,940 (10 μM) alone, in the presence of UVI3502 (10 μM) alone and in the presence of both CP55,940 and UVI3502 (both at 10 μM).

| | Stimulation (% over basal) | | | | | |
|---|---|---|---|---|---|---|
| | Sprague-Dawley rats | | | Swiss mice | | |
| **Brain region** | **CP55,940** | **UVI3502** | **CP55,940 + UVI3502** | **CP55,940** | **UVI3502** | **CP55,940 + UVI3502** |
| Amygdala | 49 ± 12 | -35 ± 19 | -47 ± 30## | 88 ± 60 | -16 ± 51* | 0 ± 78 |
| Cerebellum (gray matter) | 68 ± 77 | -16 ± 51 | -46 ± 31# | 128 ± 59 | -43 ± 9** | 6 ± 23## |
| Cortex | 40 ± 25 | -45 ± 24** | -60 ± 10## | 40 ± 31 | -55 ± 9** | -51 ± 11## |
| *Globus pallidus* | 833 ± 352 | -3 ± 46** | 89 ± 42## | 412 ± 189 | -46 ± 29* | 91 ± 104# |
| Hippocampus dorsal | 31 ± 9 | -40 ± 26* | -62 ± 6# | 77 ± 23 | -38 ± 12** | -36 ± 8# |
| *Nucleus basalis magnocellularis* | 164 ± 35 | -35 ± 30** | -40 ± 11## | 91 ± 68 | -20 ± 65* | -30 ± 37# |
| Striatum | 83 ± 81 | -34 ± 25* | -54 ± 23## | 56 ± 36 | -52 ± 18** | -52 ± 13## |
| *Substantia nigra* | 314 ± 222 | -41 ± 24** | 14 ± 80# | 747 ± 164 | -58 ± 11** | 181 ± 128# |

CP55,940 *vs*. UVI3502 (*), CP55,940 *vs*. CP55,940 + UVI3502 (#). Kruskal–Wallis test, *post-hoc* test Dunn's multiple comparison, *#p < 0.05; **##p < 0.01.
